# Niche-selective inhibition of pathogenic Th17 cells by targeting metabolic redundancy

**DOI:** 10.1101/857961

**Authors:** Lin Wu, Kate E.R. Hollinshead, Yuhan Hao, Christy Au, Lina Kroehling, Charles Ng, Woan-Yu Lin, Dayi Li, Hernandez Moura Silva, Jong Shin, Juan J. Lafaille, Richard Possemato, Michael E. Pacold, Thales Papagiannakopoulos, Alec C. Kimmelman, Rahul Satija, Dan R. Littman

## Abstract

Targeting glycolysis has been considered therapeutically intractable owing to its essential housekeeping role. However, the context-dependent requirement for individual glycolytic steps has not been fully explored. We show that CRISPR-mediated targeting of glycolysis in T cells in mice results in global loss of Th17 cells, whereas deficiency of the glycolytic enzyme glucose phosphate isomerase (Gpi1) selectively eliminates inflammatory encephalitogenic and colitogenic Th17 cells, without substantially affecting homeostatic microbiota-specific Th17 cells. In homeostatic Th17 cells, partial blockade of glycolysis upon *Gpi1* inactivation was compensated by pentose phosphate pathway flux and increased mitochondrial respiration. In contrast, inflammatory Th17 cells experience a hypoxic microenvironment known to limit mitochondrial respiration, which is incompatible with loss of *Gpi1*. Our study suggests that inhibiting glycolysis by targeting Gpi1 could be an effective therapeutic strategy with minimum toxicity for Th17-mediated autoimmune diseases, and, more generally, that metabolic redundancies can be exploited for selective targeting of disease processes.

## Introduction

Cellular metabolism is a dynamic process that supports all aspects of the cell’s activities. It is orchestrated by more than 2000 metabolic enzymes organized into pathways that are specialized for processing and producing distinct metabolites. Multiple metabolites are shared by different pathways, serving as nodes in a complex network with many redundant elements. A well-appreciated example of metabolic plasticity is the generation of ATP by either glycolysis or by mitochondrial oxidative phosphorylation (OXPHOS), making the two processes partially redundant despite having other non-redundant functions (O’Neill et al., 2016). Here, we explore metabolic plasticity in the context of T cell function and microenvironment, and the consequence of limiting that plasticity by inhibiting specific metabolic enzymes.

Th17 cells are a subset of CD4^+^ T cells whose differentiation is governed by the transcription factor RORγt, which regulates the expression of the signature IL-17 cytokines (Ivanov et al., 2006). Under homeostatic conditions, Th17 cells typically reside at mucosal surfaces where they provide protection from pathogenic bacteria and fungi and also regulate the composition of the microbiota (Kumar et al., 2016; Mao et al., 2018; Milner and Holland, 2013; Okada et al., 2015; Puel et al., 2011). However, under conditions that favor inflammatory processes, Th17 cells can adopt a pro-inflammatory program that promotes autoimmune diseases, including inflammatory bowel disease (Hue et al., 2006; Kullberg et al., 2006), psoriasis (Piskin et al., 2006; Zheng et al., 2007) and diverse forms of arthritis (Hirota et al., 2007; Murphy et al., 2003; Nakae et al., 2003a; Nakae et al., 2003b). This program is dependent on IL-23 and is typically marked by the additional expression of T-bet and IFN-*γ* (Ahern et al., 2010; Cua et al., 2003; Hirota et al., 2011; Morrison et al., 2013). The Th17 pathway has been targeted with neutralizing antibodies specific for IL-17A or IL-23 to effectively treat psoriasis, ulcerative colitis, Crohn’s disease, and some forms of arthritis (Fotiadou et al., 2018; Hanzel and D’Haens, 2020; Langley et al., 2018; Moschen et al., 2019; Pariser et al., 2018; Tahir, 2018; Wang et al., 2017), but these therapies inevitably expose patients to potential fungal and bacterial infections, as they also inhibit homeostatic Th17 cell functions. In this study, we aimed to identify genes and pathways that are required for the function of inflammatory pathogenic Th17 cells, but are dispensable for homeostatic Th17 cells, which may enable the selective therapeutic targeting of pathogenic Th17 cells.

T cell activation leads to extensive clonal expansion, demanding a large amount of energy and biomass production (O’Neill et al., 2016; Olenchock et al., 2017). Glycolysis is central both in generating ATP and providing building blocks for macromolecular biosynthesis (Zhu and Thompson, 2019). Glycolysis catabolizes glucose into pyruvate through ten enzymatic reactions. In normoxic environments, pyruvate is typically transported into the mitochondria to be processed by the tricarboxylic acid cycle, driving OXPHOS for ATP production. In hypoxic environments, OXPHOS is suppressed and glycolysis is enhanced, generating lactate from pyruvate in order to regenerate nicotinamide adenine dinucleotide (NAD^+^) needed to support ongoing glycolytic flux (Birsoy et al., 2015; Semenza, 2014; Sullivan et al., 2015). In highly proliferating cells, such as cancer cells and activated T cells, pyruvate can be converted into lactate in the presence of oxygen, a phenomenon termed aerobic glycolysis, or the “Warburg effect” (MacIver et al., 2013; Vander Heiden et al., 2009; Vander Heiden and DeBerardinis, 2017; Warburg et al., 1958; Zhu and Thompson, 2019). Although inhibiting glycolysis is an effective method of blocking T cell activation (Macintyre et al., 2014; Peng et al., 2016; Shi et al., 2011), inhibitors such as 2-Deoxy-D-glucose (2DG) have limited application in patients, due to toxic side effects as a result of the housekeeping function of glycolysis in multiple cell types (Raez et al., 2013).

Here, we systematically interrogate the requirement for individual glycolytic reactions in Th17 cell models of inflammation and homeostatic function. We found that the glycolysis gene *Gpi1* is unique in that it is selectively required by inflammatory pathogenic Th17 but not homeostatic Th17 cells. All other tested glycolysis genes were essential for functions of both Th17 cell types. Our mechanistic study revealed that upon Gpi1 loss during homeostasis, pentose phosphate pathway (PPP) activity was sufficient to maintain basal glycolytic flux for biomass generation, while increased mitochondrial respiration compensated for reduced glycolytic flux.

In contrast, Th17 cells in the inflammation models are confined to hypoxic environments and hence were unable to increase respiration to compensate for reduced glycolytic flux. This metabolic stress led to the selective elimination of Gpi1-deficient Th17 cells in inflammatory settings but not in healthy intestinal lamina propria. Overall, our study reveals a context-dependent metabolic requirement that can be targeted to eliminate pathogenic Th17 cells. As Gpi1 blockade can be better tolerated than inhibition of other glycolysis components, it is a potentially attractive target for clinical use.

## Results

### Inflammatory Th17 cells have higher expression of glycolysis pathway genes than commensal bacteria-induced homeostatic Th17 cells

Commensal SFB induce the generation of homeostatic Th17 cells in the small intestine lamina propria (SILP) (Ivanov et al., 2009), whereas immunization with myelin oligodendrocyte glycoprotein peptide (MOG) in complete Freund’s adjuvant (CFA) induces pathogenic Th17 cells in the spinal cord (SC), resulting in experimental autoimmune encephalomyelitis (EAE) (Cua et al., 2003). Th17 cell pathogenicity requires expression of IL-23R (Ahern et al., 2010; McGeachy et al., 2009). To identify genes involved in the pathogenic but not the homeostatic Th17 cell program, we reconstituted Rag1^-/-^ mice with isotype-marked *Il23r*-sufficient (*Il23r^GFP/+^*) and *Il23r* knockout (*Il23r^GFP/GFP^*) bone marrow cells, and sorted CD4^+^ T cells of both genetic backgrounds from the SILP of SFB-colonized mice and the SC of EAE mice for single cell RNA sequencing (scRNAseq) analysis (Figure S1A). We identified 4 clusters of CD4^+^ T cells in the SC of EAE mice and 5 clusters of CD4^+^ T cells in ileum lamina propria of SFB-colonized mice (Figure S1B,C)). Clusters were annotated based on their signature genes, i.e.

Th17 cells (*Il17a*), Th1 cells (*Ifng*), Treg cells (*Foxp3*), and Tcf7^+^ cells (*Tcf7*) (Figure S1D,E). Importantly, two of the EAE clusters consist mostly of IL-23R-sufficient T cells (Figures S1F-H), indicating that these populations are likely pathogenic. One of them was annotated as Th17 and the other as Th1* according to earlier publications describing such cells as *Il23r-*dependent IFN-*γ* producers (Hirota et al., 2011; Okada et al., 2015), and they were validated using flow cytometry (Figure S1I). Importantly, pathway enrichment analysis revealed glycolysis as the top pathway upregulated in both EAE Th17 and Th1* compared to the SFB-induced Th17 cells (Figures S1J,K).

We selected a subset of genes that were differentially expressed in the pathogenic Th17/Th1* cells (EAE model) versus the homeostatic Th17 cells (SFB model) for pilot functional studies (Figures S1L,M). We developed a CRISPR-based T cell transfer EAE system to evaluate the roles of the selected genes in pathogenicity (Figures S2A-C; see Methods). Among 12 genes tested, only triosephosphate isomerase 1 (*Tpi1*), encoding an enzyme in the glycolysis pathway, was required for disease onset following transfer of in vitro differentiated MOG-specific 2D2 TCR transgenic T cells (Figure S2D). This result was confirmed by analyzing *Tpi1^f/f^Cd4^cre^* mice, which, unlike Tpi1^+/+^ CD4^cre^ littermates, were completely resistant to EAE induction (Figure 1A). This finding prompted us to direct our focus on the glycolysis pathway. The relatively sparse scRNAseq data detected the transcripts of only a few glycolysis pathway genes. We therefore compared bulk RNAseq data of the two major pathogenic populations (Th1*: Klrc1^+^ Foxp3^RFP-^, Th17: IL17a^GFP+^ Klrc1^-^ Foxp3^RFP-^) and the Treg cells (Foxp3^RFP+^ IL17a^GFP-^) from the spinal cord of EAE model mice and of the Th17 (IL17a^GFP+^ Foxp3^RFP-^) and Treg (IL17a^GFP-^ Foxp3^RFP+^) cells from the SILP of the SFB model (Figures S2E, F). We found that almost every glycolysis pathway gene was expressed at a higher level in the EAE model than in the SILP CD4^+^ T cells, irrespective of whether cells were pathogenic or Treg (Figure 1B), suggesting that most CD4^+^ T cells have more active glycoysis in the inflamed spinal cord microenvironment than in the SILP at homeostasis.

**Figure 1.**
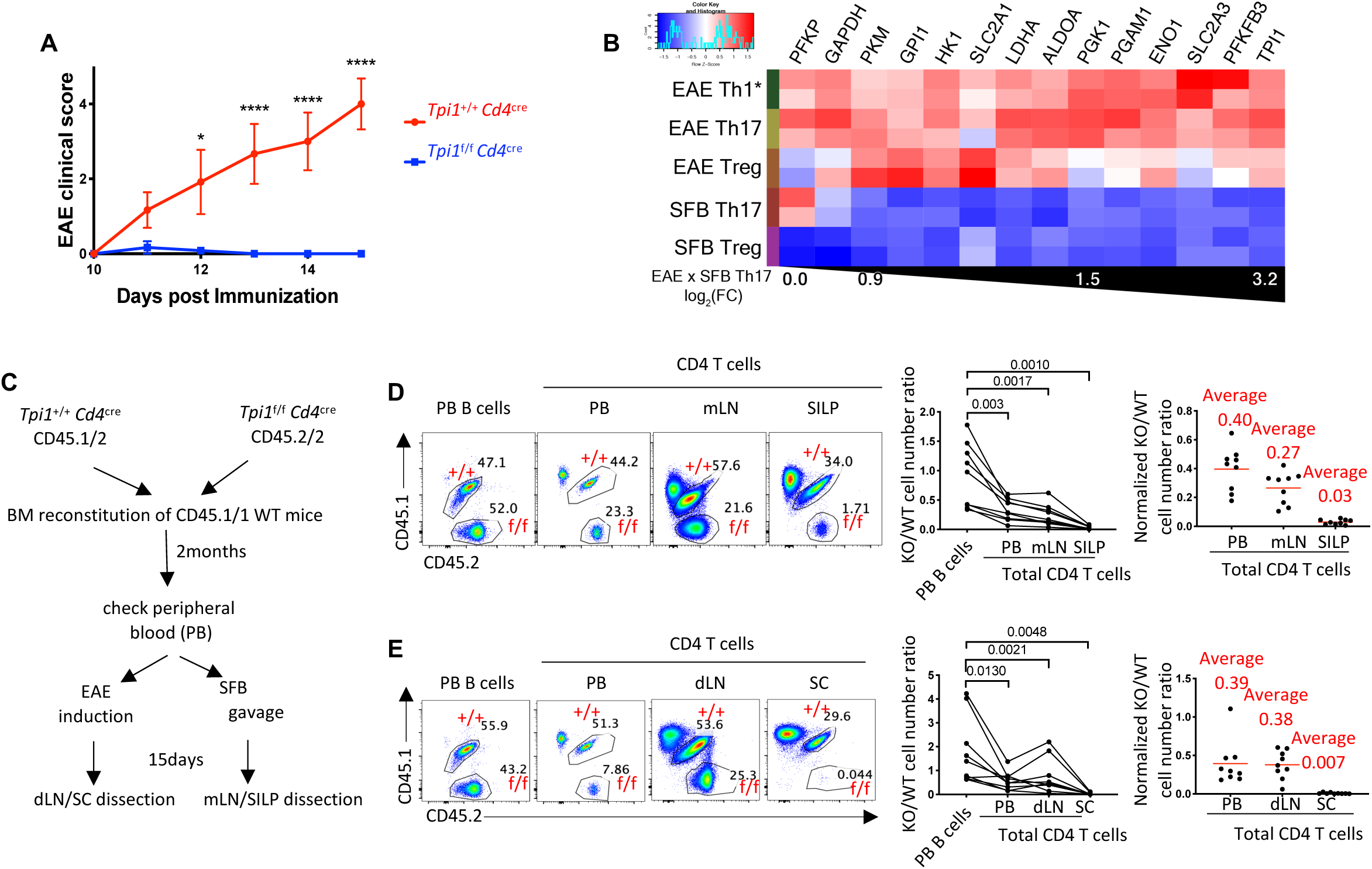
Encephalitogenic Th17 cells have higher glycolysis pathway gene expression than homeostatic SFB-induced Th17 cells. (A) Clinical disease course of MOG-CFA-induced EAE in *Tpi1^f/f^Cd4^cre^* (n=7) and *Tpi1^+/+^Cd4^cre^* littermate control mice (n=8). The experiment was repeated twice with the same conclusion. P values were determined using two-way ANOVA, * (P<0.05), **** (P<0.0001). (B) Heatmap of glycolysis pathway genes expressed in spinal cord Th1*, Th17, and Treg cells in mice with EAE (day 15, score 5) compared to Th17 and Treg cells in the SILP of SFB-colonized mice. Genes are arranged according to the fold change between EAE Th17 and SFB Th17. (C) Experimental design of the Tpi1 bone marrow reconstitution experiment. Bone marrow cells from *Tpi1*^+/+^ *Cd4*^cre^ CD45.1/2 and *Tpi1*^f/f^ *Cd4*^cre^ CD45.2/2 donor mice were transferred into lethally irradiated CD45.1/1 recipients. Peripheral blood was collected two months later for reconstitution analysis. dLN and SC of the EAE model, and mLN and SILP of the SFB model were dissected at day 15 post EAE induction or SFB colonization. (D,E) Cell number analysis of *Tpi1* WT and KO CD4^+^ T cells in the SFB model (D) and the EAE model (E). Left panels, representative FACS plots showing the frequencies of the CD45.1/2 and the CD45.2/2 B cells among all B cells in peripheral blood and of the CD45.1/2 and the CD45.2/2 CD4^+^ T cells among all CD4^+^ T cells in peripheral blood, mLN or dLN, and SILP of SFB colonized mice or SC of EAE mice. Middle panels, compilation of the cell number ratio of CD45.2/2 to CD45.1/2. Right, normalized KO/WT total CD4^+^ T cell number ratio in each tissue. The cell number ratio of peripheral B cells was set as 1. P values were determined using paired t tests. See also Figures S1,2,3.

Despite these differences in gene expression, Tpi1 was also indispensable for the SFB-dependent induction of homeostatic Th17 cells. We analyzed CD4^+^ T cell profiles in chimeric *Rag1* KO mice reconstituted with a 1:1 mix of bone marrow cells from *Tpi1^+/+^CD4^cre^* CD45.1/2 and *Tpi1^f/f^CD4^cre^* CD45.2/2 donors (Figure S3A). Tpi1 deficiency led to 33-fold reduction of total CD4^+^ T cells in the SILP and to 1.4-fold and 1.7-fold reduction in the mLN and the peripheral blood, respectively (Figure 1D). Moreover, RORγt, Foxp3, IL-17a, and IFN-*γ* were all severely reduced in mutant T cells relative to the WT cells in the SILP and mLN (Figures S3A,B). The same pattern was observed in the EAE model, using mixed bone marrow chimeric mice (Figures 1E, and S3C), indicating that Tpi1 is essential for the differentiation of all CD4^+^ T cell subsets.

### Gpi1 is selectively required by inflammatory but not homeostatic Th17 cells

The higher glycolytic activity of the inflammatory Th17 cells suggested that they may be more sensitive than the homeostatic Th17 cells to inhibition of glycolysis, which might be achieved by inactivation of *Tpi1, Gpi1 or Ldha* (explained in Figure S4A). However, neither cell type is likely to survive a complete block in glycolysis by *Gapdh* knockout. To test the hypothesis, we assessed the ability of co-tranferred control and targeted TCR transgenic T cells to differentiate into Th17 cells in models of homeostasis (7B8 T cells in SFB-colonized mice (Yang et al., 2014)) and inflammatory disease (2D2 T cells for EAE (Bettelli et al., 2003) and HH7.2 T cells for *Helicobacter*-dependent transfer colitis (Xu et al., 2018)) (Figure 2A). The genes of interest (indicated in Figure S4A) or the control gene *Olfr2* were inactivated by guide RNA electroporation of naïve isotype-marked CD4^+^ TCR and Cas9 transgenic T cells (Platt et al., 2014), which were then transferred in equal numbers into recipient mice (Figures 2A, S4B).

**Figure 2.**
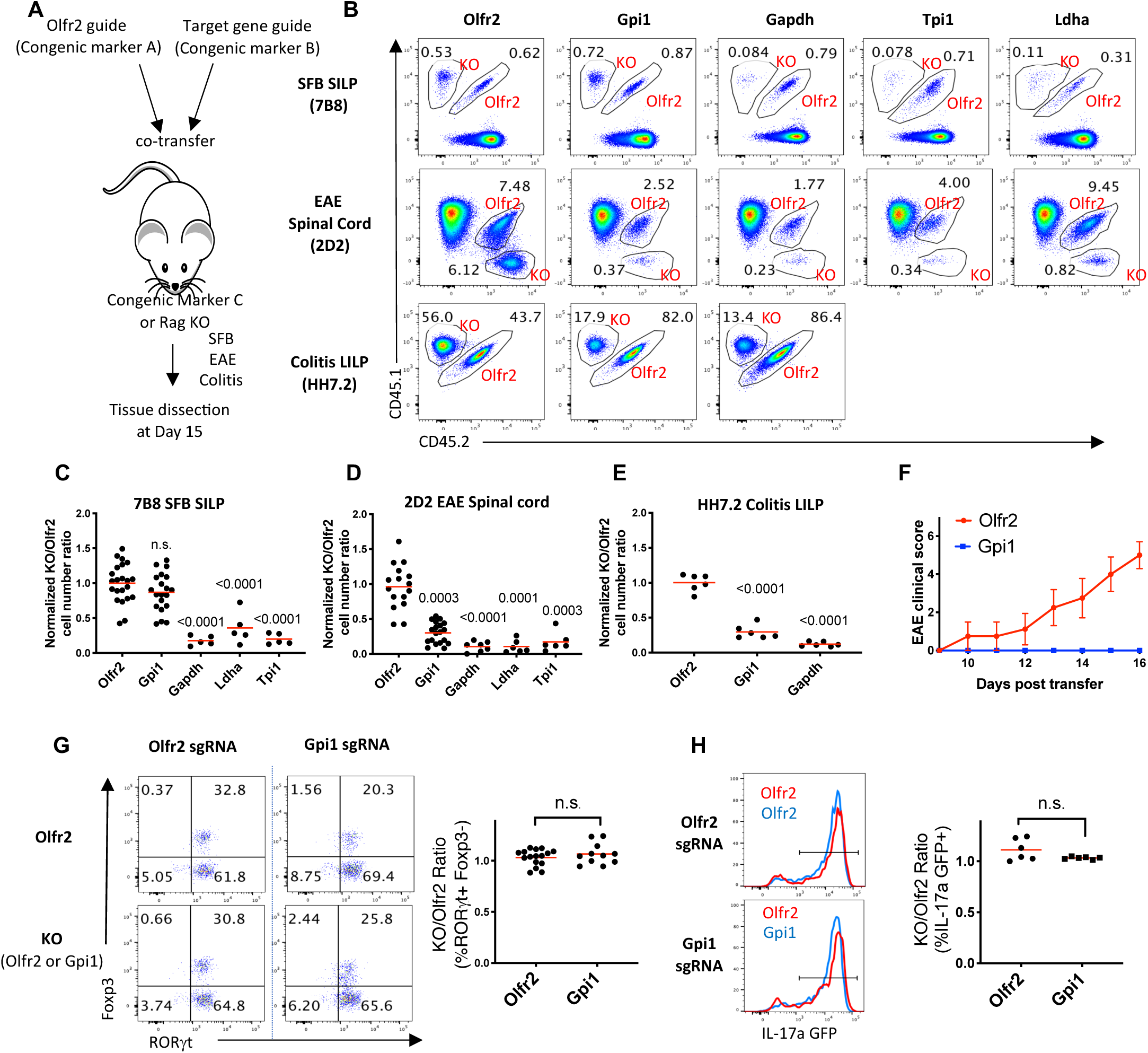
Gpi1 is selectively required by inflammatory encephalitogenic or colitogenic Th17 cells, but not by homeostatic SFB-induced Th17 cells. (A) Experimental setup. TCR transgenic Cas9-expressing naïve CD4^+^ T cells were electroporated with guide amplicons, and co-transferred with *Olfr2* amplicon-electroporated control cells into recipient mice that were immunized for EAE induction (EAE model), or had been colonized with SFB (SFB model) or *Helicobacter hepaticus* (Colitis model). (B) Representative FACS plots showing the frequencies of *Olfr2* KO cells and the co-transferred glycolytic gene KO cells among total CD4^+^ T cells in each model. (C-E) Compilation of cell number ratio of gene KO group to the co-transferred *Olfr2* control for each model, as shown in panel B. The ratio was normalized to *Olfr2*/*Olfr2* co-transfer (*Olfr2*/*Olfr2* cell number ratio was set to 1). Three independent experiments were performed with the same conclusion. (F) Clinical disease course of EAE in *Rag1* KO mice receiving *Olfr2* KO 2D2 cells or *Gpi1* KO 2D2 cells. Experiment was conducted as illustrated in Figure S2C. n=5 mice/group. The experiment was repeated twice with the same conclusion. (G) Left, representative FACS plot showing ROR*γ*t and Foxp3 expression of co-transferred targeted 7B8 cells isolated from the SILP of recipient SFB-colonized mice at day15. Right, ratio of the percentage of ROR*γ*t ^+^ Foxp3^-^ in *Gpi1*- or *Olfr2*-targeted versus co-transferred *Olfr2* control cells in each recipient. (H) Left, representative FACS histogram showing the overlay of IL17a-GFP expression of co-transferred *Olfr2* KO and *Gpi1* KO 7B8 cells isolated from the SILP of SFB-colonized mice at day15. Right, ratio of the percentage of IL-17a^+^ cells in *Gpi1*- or *Olfr2*-targeted versus co-transferred *Olfr2* control cells in each recipient. P values were determined using t tests. See also Figures S4,5.

Targeting was performed with sgRNAs that achieved the highest in vitro knockout efficiency (∼80-90%) (Figures S4C-E), which corresponded to the frequency of targeted cells at 4 days after electroporation and transfer into mice, as assessed by targeting of GFP (Figure S4B). This strategy also allowed for efficient double gene KO (Figure S4F).

In both SFB and EAE T cell transfer experiments, there were roughly 9-fold fewer *Tpi1*-targeted than co-transferred control Th17 cells in the SILP or the SC, respectively (Figures 2B-D), consistent with the *in vitro* KO efficiency (Figure S4C), suggesting that the majority of the *Tpi1* KO cells were eliminated in the total T cell pool. The remaining cells expressed RORγt and IL-17a at a similar level to control populations (Figures S5A-D), which was not observed in the previous *Tpi1* bone marrow-reconstituted mice (Figures S3B), suggesting that they were not targeted. These results indicated that the electroporation transfer system could be successfully used to evaluate T cell gene functions *in vivo*.

Similar to the *Tpi1* KO, cells deficient for *Gapdh* were eliminated in all three models tested (Figures 2B-E). Inactivation of *Ldha* resulted in loss of the cells in the EAE model, and a less striking, but still substantial, cell number reduction in the homeostatic SFB model (Figures 2B-E). In contrast, although *Gpi1* ablation reduced cell number by 75% in the inflammatory EAE and colitis models, it had no significant effect on cell number in the homeostatic Th17 cell model (Figures 2B-E). Consistent with the reduction in cell number, *Gpi1*-mutant 2D2 cells were unable to induce EAE (Figure 2F), suggesting that Gpi1 is essential for pathogenicity. We also investigated Th17 cell differentiation and cytokine production by *Gpi1*-deficient cells in the SFB model, and found no difference from co-transferred control cells in terms of RORγt/Foxp3 expression (Figures 2G) or IL-17a production, as demonstrated using 7B8 cells from mice bred to the *Il17a^GFP/+^* reporter strain (Figure 2H). *In vitro* restimulation further confirmed that transferred Gpi1-targeted 7B8 cells were indeed Gpi1 deficient (Figures S5C, D). Collectively, these data suggest that Gpi1 is selectively required by the inflammatory encephalitogenic and colitogenic Th17 cells, but not by homeostatic SFB-induced Th17 cells. In contrast, Gapdh, Tpi1 and Ldha are required for the accumulation of both types of Th17 cells.

### PPP compensates for Gpi1 deficiency in the homeostatic SFB model

To explain why Gpi1 is dispensable in the homeostatic SFB model while other glycolysis genes are not, we hypothesized that the PPP shunt might maintain some glycolytic activity in the *Gpi1* KO cells and thus compensate for Gpi1 deficiency (Figure S4A). We tested the hypothesis with *in vitro* isotope tracing. Irrespective of the non-pathogenic (npTh17, cultured in IL-6 and TGF-*β* (Veldhoen et al., 2006)) or pathogenic (pTh17, cultured in IL-6, IL-1*β*, and IL-23 (Ghoreschi et al., 2010)) conditions used, *in vitro* differentiated Th17 cells displayed less of a growth defect (25% reduction) following targeting of *Gpi1* than upon inactivation of *Gapdh* (70% reduction) (Figure 3A). This result resembles the different sensitivities observed in the homeostatic SFB model, suggesting that the cytokine milieu of the pathogenic and homeostatic Th17 cells does not account for the phenotypes of the *Gpi1* mutant mice. We incubated np/p Th17 cells with ^13^C1,2-glucose and analyzed downstream ^13^C label incorporation. The ^13^C1,2-glucose tracer is catabolized through glycolysis via Gpi1, producing pyruvate or lactate containing two heavy carbons (M2), while Gpi1-independent shunting through the PPP produces pyruvate and lactate containing one heavy carbon (M1) (Figure 3B) (Lee et al., 1998). Consistent with our hypothesis, we observed ∼ 3-fold increase in relative PPP activity in the *Gpi1* KO Th17 cells compared to the *Olfr2* KO control, irrespective of npTh17 or pTh17 condition (Figure 3C), indicating that *Gpi1* deficiency maintains active glycolytic flux through PPP shunting.

**Figure 3.**
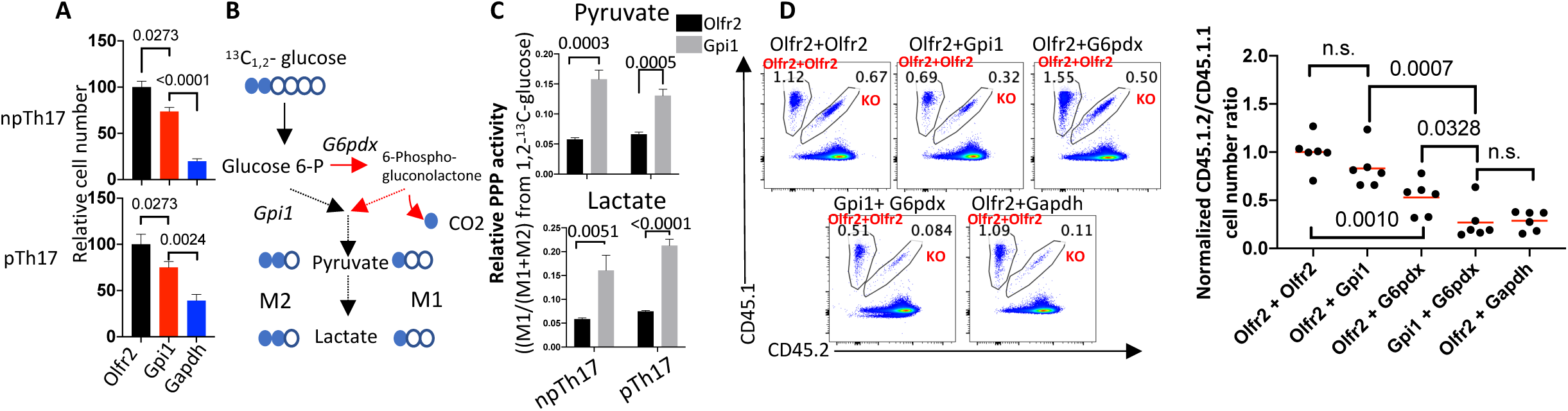
PPP activity rescues *Gpi1* deficiency in the homeostatic SFB-induced Th17 cells. (A) Relative number of Th17 cells cultured for 120 h *in vitro*. Naïve Cas9-expressing CD4^+^ T cells were electroporated with corresponding guide-amplicons and cultured in both npTh17 condition and pTh17 condition. Cell number was normalized to *Olfr2* KO group (*Olfr2* cell number was set as 100). N=3, data are representative of three independent experiments. (B) Schematic of ^13^C1,2-glucose labeling into downstream metabolites. ^13^C is labeled as filled circle. Catalytic reactions of PPP are labeled as red arrows, while those of glycolysis are in black. (C) Relative PPP activity of *in vitro* cultured Th17 cells. Relative PPP activity from ^13^C1,2-glucose was determined using the following equation: PPP= M1/ (M1+M2), where M1 is the fraction of ^13^C1,2-glucose derived from the PPP and M2 is the fraction of ^13^C1,2-glucose derived from glycolysis. (D) 7B8 Cas9 transgenic naïve CD4^+^ T cells were electroporated with a mixture of two guide amplicons (for double KO) before being transferred into recipient mice. Left, representative FACS plots showing the frequencies of co-transferred *Olfr2+Olfr2* control CD4^+^ T cells and the indicated gene KO CD4^+^ T cells in the SILP of the SFB model. Right, compilation of cell number ratios of the indicated gene KO combinations to the co-transferred *Olfr2+Olfr2* control. The ratio was normalized to *Olfr2+Olfr2*/*Olfr2+Olfr2* co-transfer. Two independent experiments were performed with same conclusion. P values were determined using t tests.

To test whether PPP activity compensates for Gpi1 deficiency *in vivo* in the homeostatic SFB model, we knocked out both *Gpi1* and *G6pdx*, which catalyzes the initial oxidative step in the PPP, with the CRISPR-electroporation transfer system. Combined deletion of these two genes would be expected to block glycolysis, similar to *Gapdh* KO. Consistent with this hypothesis, the *Gpi1/G6pdx* double KO reduced cell number by about 75%, which is similar to that with the *Gapdh* single KO, yet significantly different from that with inactivation of either *Gpi1* or *G6pdx* alone (Figure 3D). These results suggest that *in vivo* PPP activity maintains viability of the homeostatic SFB-induced Th17 cells lacking Gpi1.

### Increased mitochondrial respiration additionally compensates for Gpi1 deficiency in the homeostatic SFB model

To determine the impact of Gpi1 deficiency on aerobic glycolysis activity, we quantified lactate production by GC-MS and found it to be markedly reduced in *Gpi1* KO cells (Figure 4A). A significant reduction in lactate production may suggest compromised glycolytic ATP production in *Gpi1* KO cells. As *in vitro* cultured *Gpi1* KO Th17 cells and homeostatic SFB-specific *Gpi1* KO Th17 cells were largely unaffected in terms of cell number and cytokine production, we speculate that mitochondrial respiration may provide an alternative source of ATP to compensate. Indeed, in vitro cultured *Gpi1* KO cells displayed higher ATP-linked respiration rate than *Olfr2* control cells, irrespective of npTh17 or pTh17 cell culture condition (Figure 4B,C). Furthermore, upon Antimycin A (Complex III inhibitor) treatment, *Gpi1* KO cells were decreased in cell number, similar to *Gapdh*-deficient cells (Figure 4D), consistent with increased mitochondrial respiration compensating for Gpi1 deficiency in the *in vitro* cell culture system.

**Figure 4.**
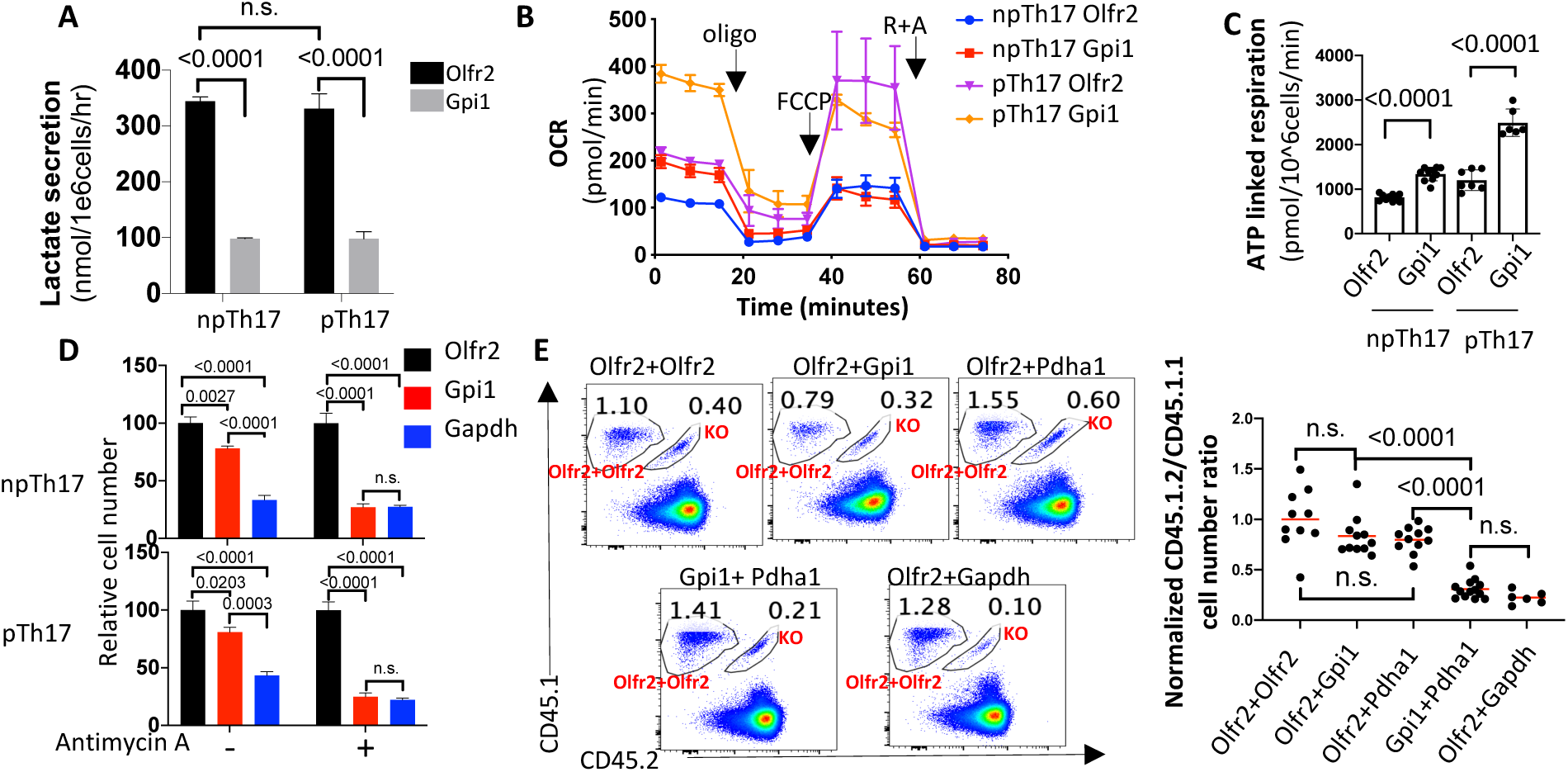
Mitochondrial respiration compensates for Gpi1 deficiency in the homeostatic SFB Th17 cells. (A) Lactate secretion of *in vitro* cultured *Olfr2* and *Gpi1* KO np/p Th17 cells. Cells were cultured as in Figure 3A. At 96 h cells were re-plated in fresh RPMI medium with 10% dialyzed FCS, and supernatants were collected 12 h later for lactate quantification by GC-MS. (B) Seahorse experiment showing the oxygen consumption rate (OCR) of *in vitro* cultured Th17 cells at baseline and in response to oligomycin (Oligo), carbonyl cyanide 4-(trifluoromethoxy) phenylhydrazone (FCCP), and rotenone plus antimycin (R + A). 10 replicates were used for the npTh17 *Olfr2* and *Gpi1* targeted cells, 7 replicates for the pTh17 *Olfr2*, and 6 replicates for the pTh17 *Gpi1* sets of targeted cells. (C) ATP linked respiration rate quantified based on experiments in panel (B). Representative data from three independent experiments. (D) Relative number of Th17 cells cultured for 120 h *in vitro* (as in Figure 3A), in the presence or absence of 10 nM antimycin A. Cell number was normalized to control (*Olfr2*-targeted cell number was set as 100). n=3, data are representative of two independent experiments. (E) 7B8 Cas9 naïve CD4^+^ T cells were electroporated with a mixture of two guide amplicons (for double KO) before transfer into recipient mice. Left, representative FACS plots showing the frequencies of co-transferred *Olfr2+Olfr2* control and indicated gene KO CD4^+^ T cells in the SILP of the SFB model at day 15. Right, compilation of cell number ratios of glycolysis gene KO group to the co-transferred *Olfr2+Olfr2* control. The ratio was normalized to *Olfr2+Olfr2*/*Olfr2+Olfr2* co-transfer. Three independent experiments were performed with same conclusion. P values were determined using t tests.

To determine whether increased respiration rescues Gpi1 deficiency *in vivo* in the homeostatic SFB model, we targeted *Gpi1* in 7B8 T cells along with genes responsible for the oxidation of three major substrates feeding the TCA cycle: *Pdha1* (pyruvate), *Gls1* (glutamine), and *Hadhb* (fatty acids). Following co-transfer of control cells and cells lacking either *Pdha1*, *Gls1*, or *Hadhb*, there was no defect in Th17 cell number, RORγt expression, or cytokine production (data not shown). However, only double knockout of *Gpi1* and *Pdha1* led to a significant reduction in cell number, similar to that with the *Gapdh* KO cells (Figure 4E), suggesting that mitochondrial respiration through pyruvate oxidation compensates for Gpi1 deficiency in homeostatic Th17 cell differentiation induced by SFB colonization.

### PPP utilization favors biosynthetic metabolite synthesis in Gpi1-deficient cells

Compensation for *Gpi1* deletion by the combination of the PPP and mitochondrial respiration suggests the existence of a partial metabolic redundancy for Gpi1. To understand the compensatory mechanism, we performed kinetic flux profiling, employing GC-MS to measure the passage of isotope label from ^13^C6-glucose into downstream metabolites (Yuan et al., 2008). The resulting kinetic data, along with metabolite abundance, was used to quantify metabolic flux in npTh17 cells prepared from *Gpi1* KO, *Olfr2* KO and cells treated with koningic acid (KA), to inhibit Gapdh (Liberti et al., 2017), since more than 50% of cells in the *Gapdh* KO culture were WT escapees (Figure S4C)).

Loss of Gpi1 resulted in reduction of glucose uptake (Figure 5A) and in substantially decreased transmission of isotopic label to intracellular pyruvate and lactate, whose abundance was also reduced (Figures 5C,D,I). Interestingly, a lower lactate production/glucose consumption ratio was observed in *Gpi1* KO than control cells (Figure 5B), suggesting that Gpi1-deficient cells may be more inclined to use glucose carbons for the synthesis of biosynthetic metabolites, rather than lactate production. Indeed, the labeling rate and total abundance of the biosynthetic metabolites alanine, serine, glycine, and citrate either did not change or were slightly increased in the *Gpi1* KO cells (Figures 5E-I). The flux of pyruvate, lactate, alanine, serine, and glycine in the Gpi1-deficient cells, as compared to *Olfr2* KO control (Figure 5J), was consistent with the PPP supporting the branching pathways of glycolysis important for biosynthetic metabolite production, despite a reduced glycolysis rate. In contrast, Gapdh inhibition by KA treatment almost completely prevented the synthesis of downstream metabolites (Figures 5C-H) and lactate production (Figure 5K). However, it did not induce an increase in OCR (Figures 5K, L), possibly due to the severely limited production of pyruvate.

**Figure 5.**
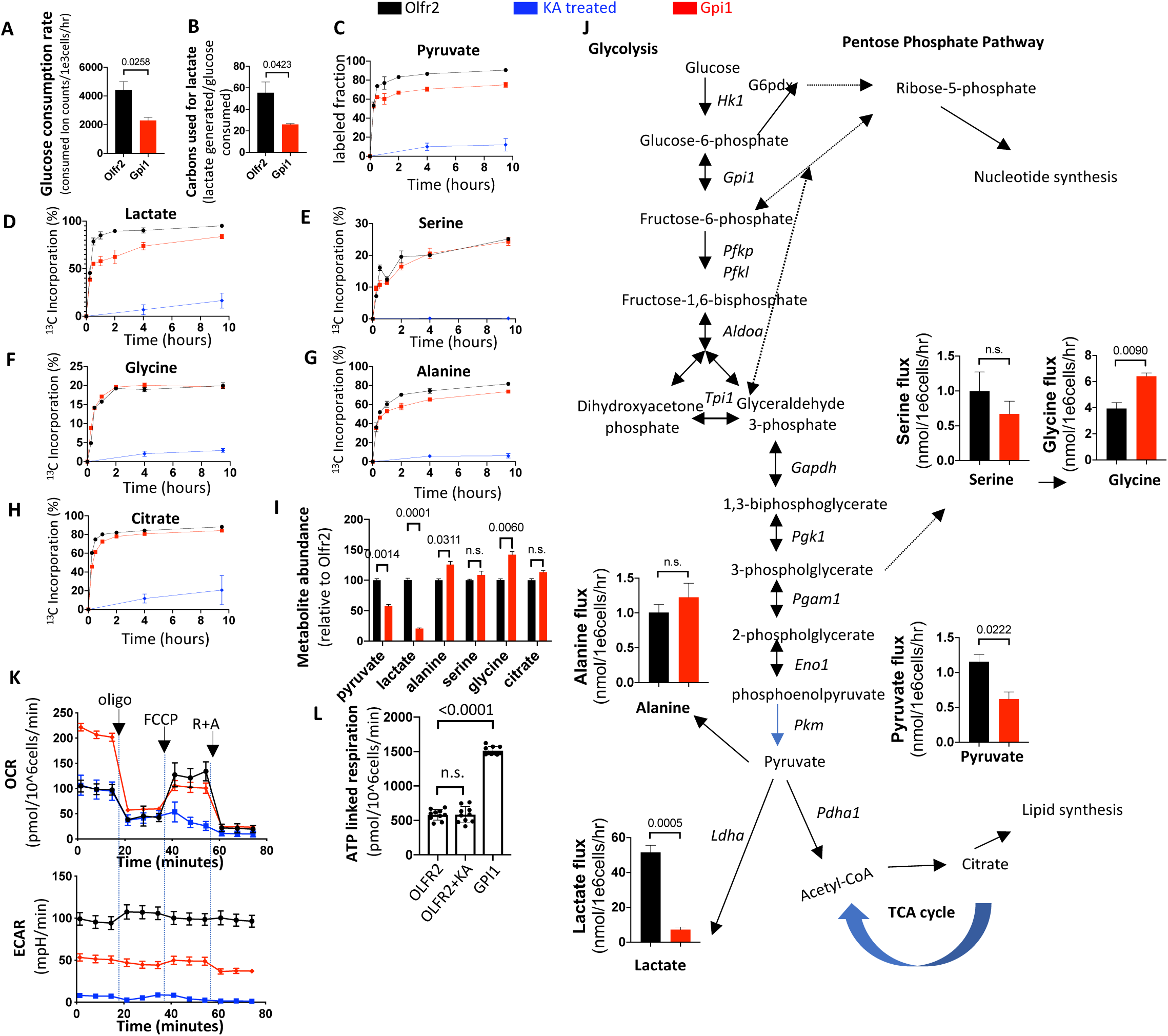
PPP supports normal biomass synthesis in the *Gpi1*-deficient npTh17 cells. *Olfr2-* or *Gpi1-*targeted npTh17 cells (cultured for 96 h as in Figure 3A) were collected for further metabolic analysis. (A, B) Cells were re-plated in U-^13^C6-glucose tracing medium to measure (A) glucose consumption rate, and (B) ratio of released lactate to consumed glucose by LC-MS. (C-H) Cells were re-plated in U-^13^C6-glucose tracing medium, and cell pellets were collected at different time points to measure ^13^C incorporation kinetics of pyruvate (C), lactate (D), serine (E), glycine (F), alanine (G) and citrate (H) in *Olfr2* KO, *Gpi1* KO, or koningic acid (KA)-treated npTh17 cells by GC-MS. (I) Intracellular abundance of pyruvate, lactate, alanine, serine, glycine and citrate in targeted npTh17 cells, as determined by GC-MS. Raw ion counts were normalized to extraction efficiency and cell number. The abundance is shown relative to the *Olfr2* sample set as 100. (J) Quantification of the production flux of serine, glycine, alanine, pyruvate and lactate based on the ^13^C incorporation kinetics and intracellular abundance of each metabolite. Three replicates for each sample. (K) OCR (top) and Extracellular Acidification Rate (ECAR) (bottom) of targeted or KA-treated npTh17 cells at baseline and in response to oligomycin (Oligo), carbonyl cyanide 4-(trifluoromethoxy) phenylhydrazone (FCCP), and rotenone plus antimycin (R + A). 10 replicates for each condition. (L) ATP-linked respiration rate, quantified based on panel K. Data are representative of two independent experiments. P values were determined using t tests.

Taken together, we conclude that the reduced amount of glucose metabolized via the PPP by *Gpi1*-deficient cells was sufficient to support the production of glycolytic intermediates and maintain pyruvate oxidation. *Gpi1*-deficient cells increased their mitochondrial respiration to compensate for the loss of glycolytic flux. In contrast, glycolytic blockade by Gapdh inhibition completely prevented carbon flux needed for key biosynthetic metabolites, which is incompatible with cell viability.

### Gpi1 is essential in the hypoxic setting of Th17-mediated inflammation

To understand why inflammatory Th17 cells are particularly sensitive to Gpi1 deficiency, we reasoned that metabolic compensation (either through PPP or mitochondrial respiration), which occurs in the homeostatic model, may be restricted in inflammatory Th17 cells. *G6pdx* KO in the EAE model resulted in a 70% reduction in 2D2 cell number in the spinal cord (Figures S6A,B), suggesting that PPP flux is still active in the setting of inflammation. On the other hand, a number of reports have shown that inflamed tissues, including the spinal cord in EAE and the LILP in colitis, are associated with low oxygenation (hypoxia) (Davies et al., 2013; Johnson et al., 2016; Karhausen et al., 2004; Naughton et al., 1993; Ng et al., 2010; Peters et al., 2004; Van Welden et al., 2017; Yang and Dunn, 2015). This suggests that impaired mitochondrial respiration might occur which would result in an inability to provide sufficient ATP to compensate for energy loss due to Gpi1 deficiency in these tissues. To test this hypothesis, we cultured Th17 cells in normoxic (20% O2) and mild hypoxic (3% O2) conditions, and found that *Gpi1* KO Th17 cells in the hypoxic state, irrespective of npTh17 or pTh17 differentiation protocols, displayed a growth defect similar to that occurring with Gapdh deficiency (Figure 6A). Furthermore, Olfr2 control npTh17 cells displayed about a 2-fold increase in lactate production in hypoxic compared to normoxic conditions (Figure S6C), indicating that hypoxic cells rely more on glycolysis to generate ATP. *Gpi1* KO cells had substantially decreased lactate production in both normoxic and hypoxic conditions (Figure S6C), but there was reduced intracellular ATP only with hypoxia (Figure S6D), suggesting that restrained mitochondrial respiration in hypoxia cannot compensate for Gpi1 inactivation, and hence leads to energy crisis.

**Figure 6.**
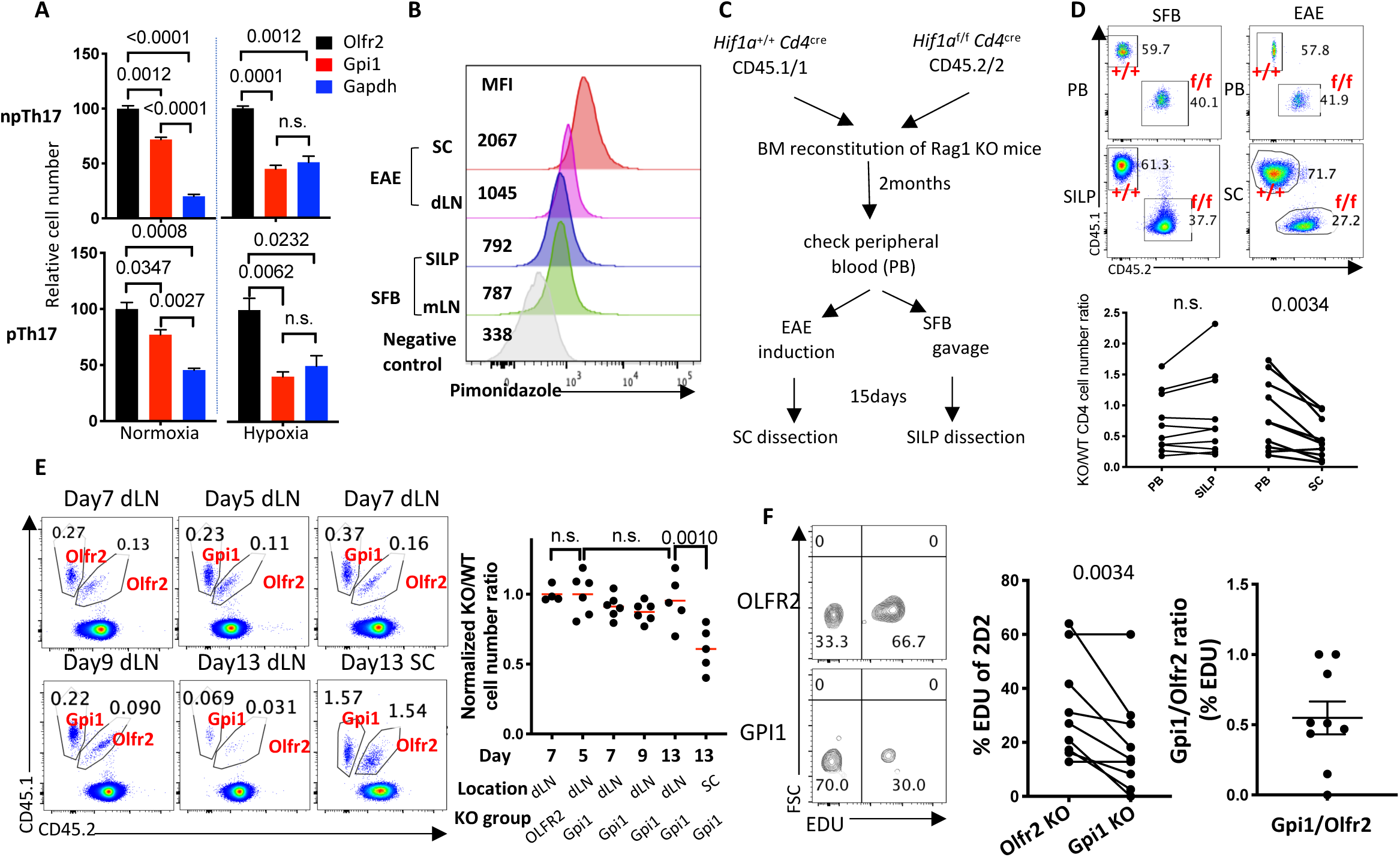
Hypoxia in the inflamed spinal cord of the EAE model restrains mitochondrial respiration, leading to the elimination of Gpi1-deficient Th17 cells. (A) Relative number of gRNA-targeted Th17 cells cultured for 120 h *in vitro* in either normoxic (20% O2) or hypoxic (3% O2) conditions. Cell number was normalized to *Olfr2* KO (*Olfr2* cell number was set as 100). (B) Representative FACS histogram showing pimonidazole labeling of total CD4^+^ T cells isolated from the SC and the dLN at day 15 after induction of EAE and from the SILP and the mLN of SFB-colonized mice. (C) Experimental design of the Hif1a bone marrow reconstitution experiment. *Rag1* KO mice were lethally irradiated and reconstituted with equal numbers of bone marrow cells from *Hif1a*^+/+^ *Cd4*^cre^ CD45.1/2 and *Hif1a*^f/f^ *Cd4*^cre^ CD45.1/1 donors. CD4^+^ T cells were examined 2-months later in peripheral blood (PB), after which mice were either gavaged with SFB or immunized for EAE induction, and CD4^+^ T cells from SC or SILP from each model were isolated at day15 for examination. (D) Top, representative FACS plots showing the frequencies of *Hif1a*^+/+^ *Cd4*^cre^ and *Hif1a*^f/f^*Cd4*^cre^ CD4^+^ T cells in the peripheral blood and the SILP or the SC of the SFB and the EAE models, respectively. Bottom, compilation of the KO/WT total CD4^+^ cell number ratio obtained in the PB and the SILP or the SC of the SFB and EAE models. Two experiments were performed with the same conclusion. (E) Time-course of 2D2 cas9 co-transfer experiment. Left, representative FACS plot showing the frequencies of the *Olfr2* control and the co-transferred *Olfr2* or *Gpi1* KO 2D2 cells in total CD4^+^ T cells isolated from the dLN or the SC at different time points of EAE. Right, compilation of the KO/Olfr2 control cell number ratio in the dLN or the SC. Data are representative of two independent experiments. (F) EDU in vivo labeling of co-transferred 2D2 cells in the EAE model. Left, representative FACS plots showing the frequencies of EDU^+^ among co-transferred *Olfr2* control and *Gpi1* KO 2D2 cells in the SC at the onset of EAE (day10). Middle, compilation of the percentage EDU^+^ cells in the *Olfr2* control and the *Gpi1* KO cells. Right, the *Gpi1* KO/*Olfr2* control ratios of percentage EDU^+^ cells. Two experiments were performed with the same conclusion. P values were determined using t tests. Panel D, F, and Day13 dLN and SC comparison in panel E were analyzed with paired t tests. See also Figure S6.

We next investigated whether the inflamed tissue in the EAE model differs from the homeostatic tissue in the SFB model with regard to oxygen availability. To address this, we performed I.V. injection of pimonidazole, an indicator of O2 levels, into mice with EAE or with SFB colonization, and found that CD4^+^ T cells in the EAE model spinal cord had two-fold higher pimonidazole staining than in the dLN and ∼2.7-fold compared to CD4^+^ T cells in SILP and mLN (Figure 6B). This result suggests that CD4^+^ T cells in the spinal cord of the EAE model experience greater oxygen deprivation than in other tissues examined. The increased labeling in the spinal cord was observed in all subsets of CD4^+^ T cells (Figures S6E,F), indicating that decreased oxygen availability is likely a tissue feature.

Hif1*α*is an oxygen sensor that, upon activation by hypoxia, initiates transcription of genes, including glycolysis genes, to allow cells to adapt in poorly oxygenated tissue (Corcoran and O’Neill, 2016; Lee et al., 2020; Semenza, 2013). To further examine the oxygenation level of SILP and SC, we investigated the effect of *Hif1a* deficiency in the different models of CD4^+^ T cell differentiation. Using chimeric *Rag1* KO mice reconstituted with equal numbers of *Hif1a^+/+^CD4^cre^* and *Hif1a^f/f^CD4^cre^* bone marrow cells, there was ∼ 40% reduction of Hif1*α*-deficient CD4^+^ T cells in the spinal cord of mice with EAE, but little difference in the SILP (Figures 6C-E). Furthermore, in the SILP of SFB-colonized mice, there was no significant difference in RORγt and Foxp3 expression or IL-17a and IFN-*γ* production between WT and KO CD4^+^ T cells (Figures S6G,H). In contrast, in the spinal cord of the EAE mice, there was a slight decrease in the Th17 cell fractions and increase in the Foxp3^+^ fraction in the *Hif1a* KO compared to WT CD4^+^ T cells (Figures S6G, H). Overall, these data suggest that Hif1*α*is functionally important for the pathogenic Th17 cells (as well as Treg cells) in the EAE model, but not for homeostatic SILP Th17 cells, reinforcing the conclusion of the pimonidazole experiment that T cells in the inflamed spinal cord experience greater oxygen deprivation than those in the healthy small intestine lamina propria.

To address when and how Gpi1 deficiency impairs inflammatory Th17 cell differentiation and/or function, we performed a time course experiment to assess the kinetics of the co-transferred *Olfr2* KO control and *Gpi1* KO 2D2 cells in the EAE model. *Gpi1* KO T cells, like *Olfr2* KO control cells, were maintained in the dLN until day 13, the last time point when transferred 2D2 cells were still detectable in the dLN (Figure 6E). However, at 13 days post immunization, *Gpi1* mutant cell number was reduced by 50% compared to control T cells in the spinal cord (Figure 6E). This finding is consistent with the previous observation that the spinal cord, but not the dLN, is hypoxic. Accordingly, we observed reduced proliferation of *Gpi1* KO 2D2 cells in the spinal cord but not in the dLN (Figures 6F and S6I). Importantly, 2D2 cells in the dLN of mice with EAE proliferated at a markedly faster rate than 7B8 cells in the mLN of the SFB model (Figures S6J,K). This demonstrates that Gpi1-deficient 2D2 cells can maintain the higher demand for energy and biomass associated with rapid proliferation and emphasizes the role of the SC hypoxic environment in the T cell’s dependence on Gpi1. We also examined Th17 cell differentiation/cytokine production, by staining for ROR*γ*t/Foxp3 and IL-17a, apoptosis, by staining for the active form of caspase 3, and potential for migration, by staining for Ccr6, and observed no differences between *Gpi1*-deficient and co-transferred control *Olfr2* KO 2D2 T cells (Figures S6L-P). We conclude that Th17 cells increase glycolysis activity to adapt to the hypoxic environment in the EAE spinal cord, and the restrained mitochondrial respiration in this setting cannot compensate for the loss of glycolytic ATP production upon Gpi1 inactivation, leading to energy crisis and cell elimination.

## Discussion

Our results demonstrate the remarkable plasticity of cellular metabolism owing to redundant components in the network, and highlight that plasticity can vary based on microenvironmental factors. Due to the unique position of Gpi1 in the glycolysis pathway, its deficiency was compensated by the combination of two metabolic pathways – the PPP and mitochondrial respiration. The PPP maintained glycolytic activity, albeit at a reduced level, to fully support biosynthetic precursor synthesis through branching pathways. Pyruvate, which is also produced by way of the PPP, increased its flux into the TCA cycle, elevating ATP production from mitochondrial respiration, thus compensating for loss of ATP production from glycolysis. Therefore, upon Gpi1 inactivation, homeostatic SFB-induced Th17 cells reprogrammed their metabolism, retaining the ability to provide adequate biomass and energy to support cell function. In contrast, limitation of mitochondrial respiration due to low oxygen availability in the inflamed EAE spinal cord rendered Gpi1-deficient cells unable to generate sufficient ATP molecules either through glycolysis or mitochondrial respiration, resulting in energy crisis and cell elimination. Inactivation of Gapdh, whose function, unlike that of Gpi1, cannot be compensated by other metabolic pathways, completely blocked glycolysis and hence led to elimination of both homeostatic and inflammatory Th17 cells. Our results suggest not only that suppressing glycolysis by Gpi1 inhibition could be a well-tolerated treatment for hypoxia-related autoimmune diseases, but also that metabolic redundancies can be exploited for targeting disease processes in selected tissues.

### Metabolic requirements for CD4^+^ T cell activation and differentiation

We found that CD4^+^ T cells (pathogenic or Treg) in the EAE model had greater expression of glycolytic genes than homeostatic CD4^+^ T cells, which is consistent with the recent demonstration that *Citrobacter*-induced inflammatory Th17 cells were more glycolytic than SFB-induced homeostatic Th17 cells (Omenetti et al., 2019). Our data suggest that T cells adapt to the hypoxic environment in the inflamed spinal cord by up-regulating glycolysis, most likely by hypoxia-stabilized Hif1*α*.

In spite of the difference in glycolytic activity, blockade of glycolysis by *Gapdh* KO is incompatible with Th17 cell expansion and differentiation in either homeostatic or inflammatory models. However, the finding that Gpi1-deficient 2D2 T cells proliferated normally in the dLN of the EAE model suggests that T cell activation and differentiation *in vivo* do not necessarily need a fully functional glycolytic pathway. With the PPP providing sufficient metabolites for biomass production and mitochondria generating greater amounts of ATP, T cell proliferation and execution of the non-pathogenic Th17 program *in vivo* can be largely maintained in the *Gpi1* KO cells. Our data highlight the complexity of the glycolysis pathway and suggest that not every enzyme is equally important to glycolytic activity and cell function.

Treg cell differentiation has been proposed to depend on OXPHOS (Kullberg et al., 2006; Macintyre et al., 2014; MacIver et al., 2013; Michalek et al., 2011), and was promoted by glycolysis inhibition *in vitro* with 2DG (Shi et al., 2011). Our *in vivo* genetic data however support an indispensable role of glycolysis in Treg cell differentiation, as demonstrated by complete loss of the intestinal and SC Foxp3^+^ T cells in mice deficient for *Tpi1* in T cells.

One intriguing finding that we have not characterized in detail is the dispensable role of Pdha1, Gls1, and Hadhb in SFB-induced homeostatic Th17 cell differentiation. While OXPHOS is essential for SFB-specific Th17 cells (data not shown), their ability to differentiate in the absence of genes controlling metabolite fueling of the TCA cycle suggests potential redundancies. Furthermore, the discrepancy between the in vivo and in vitro function of Pdha1(as shown in this study) and Gls1 (by comparing the in vivo data in this study with reported in vitro data (Johnson et al., 2018)) suggests that the microenvironment is critical for T cell metabolism and gene function, as does a recent report showing distinct metabolism profiles of in vivo and in vitro activated T cells (Ma et al., 2019).

### Gpi1 as a drug target for hypoxia-related diseases

Our study reveals that Gpi1 may be a good drug target for Th17-mediated autoimmune diseases. The sensitivity to Gpi1 deficiency is dependent on whether cells are competent to increase mitochondrial respiration. Therefore, other disease processes involving hypoxia-related pathologies may also be susceptible to Gpi1 inhibition. A recent report showed that 1% O2 almost completely inhibited *in vitro* growth of *Gpi1*-mutant tumor cell lines, but the proliferation defect was much milder when cells were cultured in normoxic condition (de Padua et al., 2017). This suggests that hypoxia-mediated sensitivity to Gpi1 deficiency can be a general phenomenon, from Th17 cells to tumor cells, which may extend the deployment of Gpi1 inhibition to a broader range of diseases.

Sensitivity to inhibition of both Hif1*α* and Gpi1 is dependent on oxygen availability. In the mixed bone marrow chimera experiments with the EAE model, there was less than a 2-fold reduction in *Hif1a* mutant compared to wild-type spinal cord CD4^+^ T cells, whereas *Gpi1*-targeted T cells were reduced 5-fold compared to wild-type cells in the transfer model of EAE. *Hif1a* inactivation efficiency was almost complete (data not shown), while *Gpi1* KO efficiency was roughly 80%, which suggests that Gpi1 deficiency resulted in a more severe defect than *Hif1a* deficiency when the T cells were in a hypoxic environment. This is not unexpected, as Gpi1 itself participates in the glycolysis pathway while Hif1*α* regulates the transcription of genes in the pathway. Although EAE was attenuated following *Hif1a* inactivation in CD4^+^ T cells (Dang et al., 2011; Shi et al., 2011), our results suggest that Gpi1 inhibition may be at least as effective a means of blocking cell function in hypoxic microenvironments.

While our data suggest that hypoxia is an essential factor that restains proliferation of Gpi1-deficient Th17 cells in the spinal cord, we cannot exclude the possibility that other environmental factors also contribute to the phenotype, as oxygenation level is certainly not the only difference between the inflamed spinal cord and the healthy small intestine lamina propria.

### Gpi1 targeting can be tolerated

Glycolysis is a universal metabolic pathway whose blockade by inhibitors such as 2DG can yield adverse side effects even at dosages too low to control tumor growth (Raez et al., 2013). 2DG inhibits the first three steps of glycolysis, an effect that is recapitulated by Gapdh inhibition, given that no alternative pathway exists for glucose catabolism. Although glycolysis is essential for T cell activation, treating diseases caused by autoimmune T cells with drugs that fully inhibit the pathway, e.g. 2DG or KA, would also likely be too toxic at potentially efficacious dosages. In light of the results presented here, Gpi1 may represent a better drug target than other glycolysis gene products for treating Th17 cell-mediated autoimmune diseases, as targeting Gpi1 may be well tolerated by most cells and tissues, thus causing minimal side effects.

This hypothesis is supported by genetic data obtained from patients with glycolysis gene deficiencies. With few exceptions, patients deficient for *Tpi1* or *Pgk1* have combined symptoms of anemia, mental retardation and muscle weakness [Online Mendelian Inheritance in Man (OMIM) entry 190450, 311800] (Schneider, 2000). Mutations in glycolysis gene products that have redundant isozymes can result in dysfunction of cell types in which only the mutated isozyme gets expressed, such that *Pgam2* deficiency results in muscle breakdown and *Pklr* deficiency causes anemia (Zanella et al., 2005). The human genetic data are thus consistent with the prediction that inhibition of glycolysis enzymes would result in serious side effects. However, the vast majority of patients with *Gpi1* mutations, which can result in preservation of less than 20% enzymatic activity, have mild to severe anemia that is treatable, with no neurological or muscle developmental defects (Baronciani et al., 1996; Kanno et al., 1996; Kugler et al., 1998; McMullin, 1999; Zaidi et al., 2017) [OMIM entry 172400]. These clinical data indicate that neurons and muscle cells can afford losing most of their Gpi1 activity even though they require glycolysis, and hence they behave similarly to homeostatic Th17 cells. Collectively, these patient data strongly support our proposal that Gpi1 can be a therapeutic target.

### Selective metabolic redundancies permit selective cell inhibition

Metabolic redundancy has been demonstrated in many biological systems, from bacteria to cancer cell lines (Bulcha et al., 2019; Deutscher et al., 2006; Horlbeck et al., 2018; Segre et al., 2005; Thiele et al., 2005; Zhao et al., 2018). Our study demonstrates compensatory metabolic pathways in CD4^+^ T cells *in vivo*, and highlights that such redundancy can be selective, depending on the cellular microenvironment. Indeed, specific inhibition of pathogenic cells can be achieved by metabolic targeting of selective redundant components. Therefore, characterization of the metabolic redundancy of cells in microenvironments with varying oxygen level, pH, abundance of extracellular metabolites, redox status, cytokine milieu, or temperature, may provide opportunities for metabolic targeting of cells in selected tissue microenvironments for therapeutic purposes.

## Acknowledgements

We thank the NYU Langone Genome Technology Center for expert library preparation and sequencing. This shared resource is partially supported by the Cancer Center Support Grant P30CA016087 at the Laura and Isaac Perlmutter Cancer Center. We thank Drew R. Jones and Rebecca E. Rose in the NYU Metabolomics Laboratory for helpful discussion and aid with LC-MS data generation and analysis. We thank S.Y. Kim in the NYU Rodent Genetic Engineering Laboratory (RGEL) for generating the Tpi1 conditional KO mice. We thank Anne R. Bresnick (Department of Biochemistry, Albert Einstein College of Medicine) for sharing the S100a4 mutant mice. This work was supported by National Multiple Sclerosis Society Fellowship FG 2089-A-1 (L.W.), by Immunology and Inflammation training grant T32AI100853 (C.N.), by NIH grant R01AI121436 (D.R.L.), by the Howard Hughes Medical Institute (D.R.L.), and the Helen and Martin Kimmel Center for Biology and Medicine (D.R.L.)

## Author Contributions

L.W. and D.R.L. conceived the project, and wrote the manuscript. L.W. designed, performed experiments, and analyzed the data. L.W. and K.E.R.H. designed, performed the GC-MS flux, tracing, and seahorse experiments, analyzed the data. Y.H. and R.S. analyzed the scRNAseq data. L.K. analyzed the bulk RNAseq data. C.A., W.L., D.L., and H.M.S. helped with mouse experiments. C.N. and W.L. helped with the development of the CRISPR/Cas9 transfection method. S.J. and R.P. helped with GC-MS experiments. K.E.R.H., J.J.L., R.P., M.E.P., T.Y.P., and A.C.K. assisted with the analysis and interpretation of the metabolic data. K.E.R.H., H.M.S., S.J., J.J.L., R.P., M.E.P., T.Y.P., A.C.K., and R.S. edited the manuscript. D.R.L. supervised the work.

## Supplemental figure titles and legends

**Figure S1.**
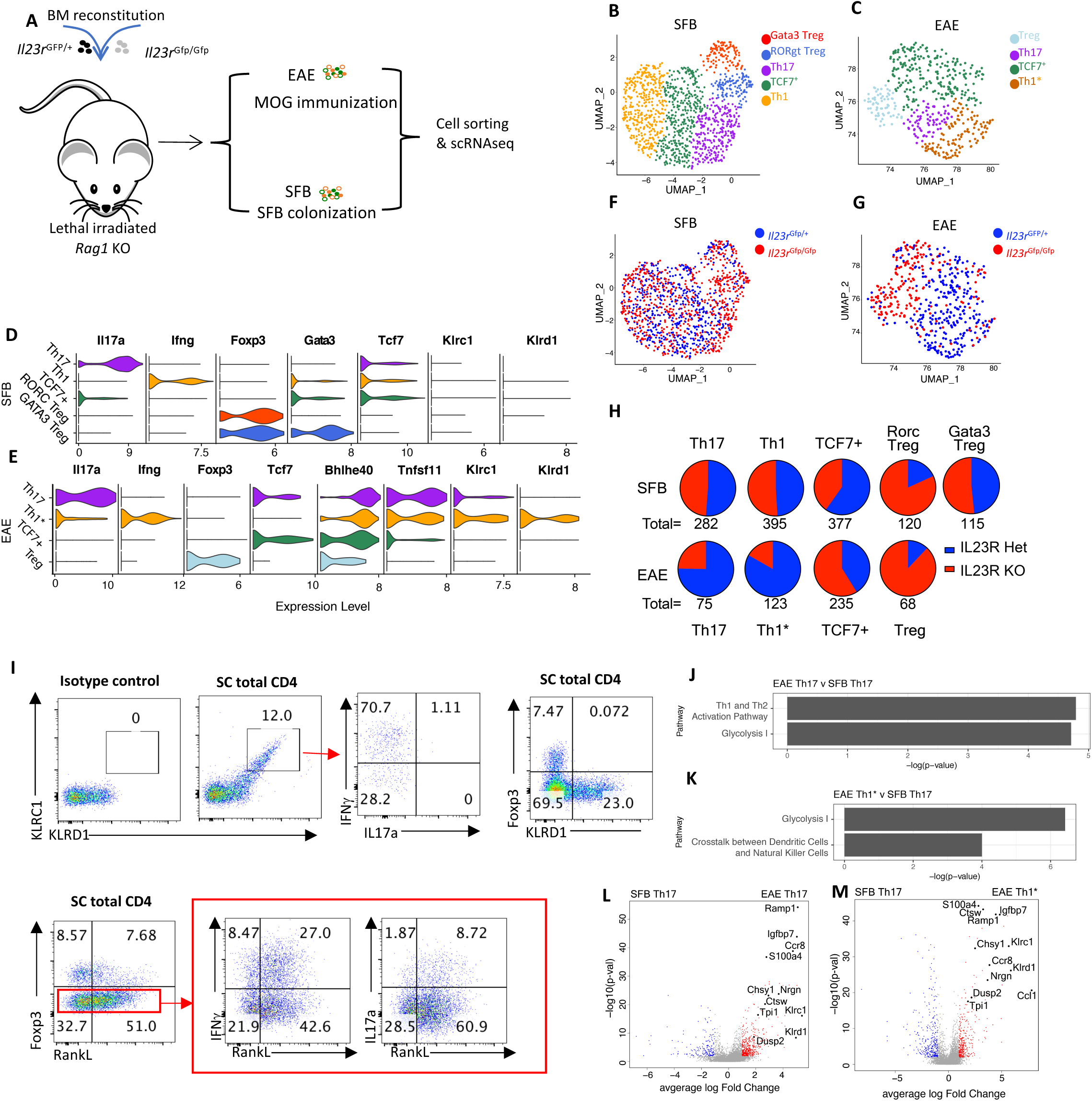
Single cell RNAseq analysis of ex vivo CD4^+^ T cells from spinal cord in the EAE model and the SILP of SFB-colonized mice, related to Figure 1. (A) Experimental design of the scRNAseq analysis. Lethally irradiated *Rag1* KO mice were reconstituted with equal numbers of bone marrow cells from isotype-marked *Il23r*^Gfp/+^ and *Il23r*^Gfp/Gfp^ donors. The CD4^+^ T cells of both *Il23r* het and KO background in the SC and SILP of each model were sorted for scRNAseq analysis. (B, C) UMAP projections of the scRNAseq data showing the five CD4^+^ T cell sub-populations in the SILP of SFB colonized mice and the four sub-populations in the SC of EAE mice. (D, E) Violin plots showing the expression level of selected marker genes for each cluster in the SFB model (D) and the EAE model (E). (F, G) Distribution of *Il23r*^Gfp/+^ and *Il23r*^Gfp/Gfp^ cells in the UMAP projection of SILP CD4^+^ T cells (F) and EAE SC CD4^+^ T cells (G). (H) Pie chart showing the percentage of *Il23r*^het^ and *Il23r*^KO^ cells in each cluster as in panels D-G. The total number of *Il23r*^Gfp/+^ cells is scaled according to that of *Il23r*^Gfp/Gfp^ cells for the analysis (assuming same number of heterozygous and homozygous cells were analyzed). The actual cell number of each cluster is indicated beneath the corresponding chart. (I)Validation of marker gene expression in EAE clusters shown in panel E. Top, Klrc1 and Klrd1 are co-expressed on cells that mainly produce IFN-*γ* (left), but are not expressed on Treg cells (right), and are thus considered as Th1* (IFN-*γ* producing pathogenic Th17) markers. Bottom left, Treg cells express less Rankl (encoded by *Tnfsf11*). Bottom right, majority of the IFN-*γ*- and IL-17a-producing cells are Rankl^+^. (J, K) Pathway enrichment analysis showing the top 2 pathways activated in EAE-associated Th17 (J) or Th1* (K) cells compared to SFB-induced Th17 cells. (L, M) Volcano plots based on the scRNAseq analysis showing the differentially expressed genes in EAE-associated Th17 (L) or Th1* cells (M) compared to SFB-induced Th17 cells. Genes selected for functional screening are highlighted.

**Figure S2.**
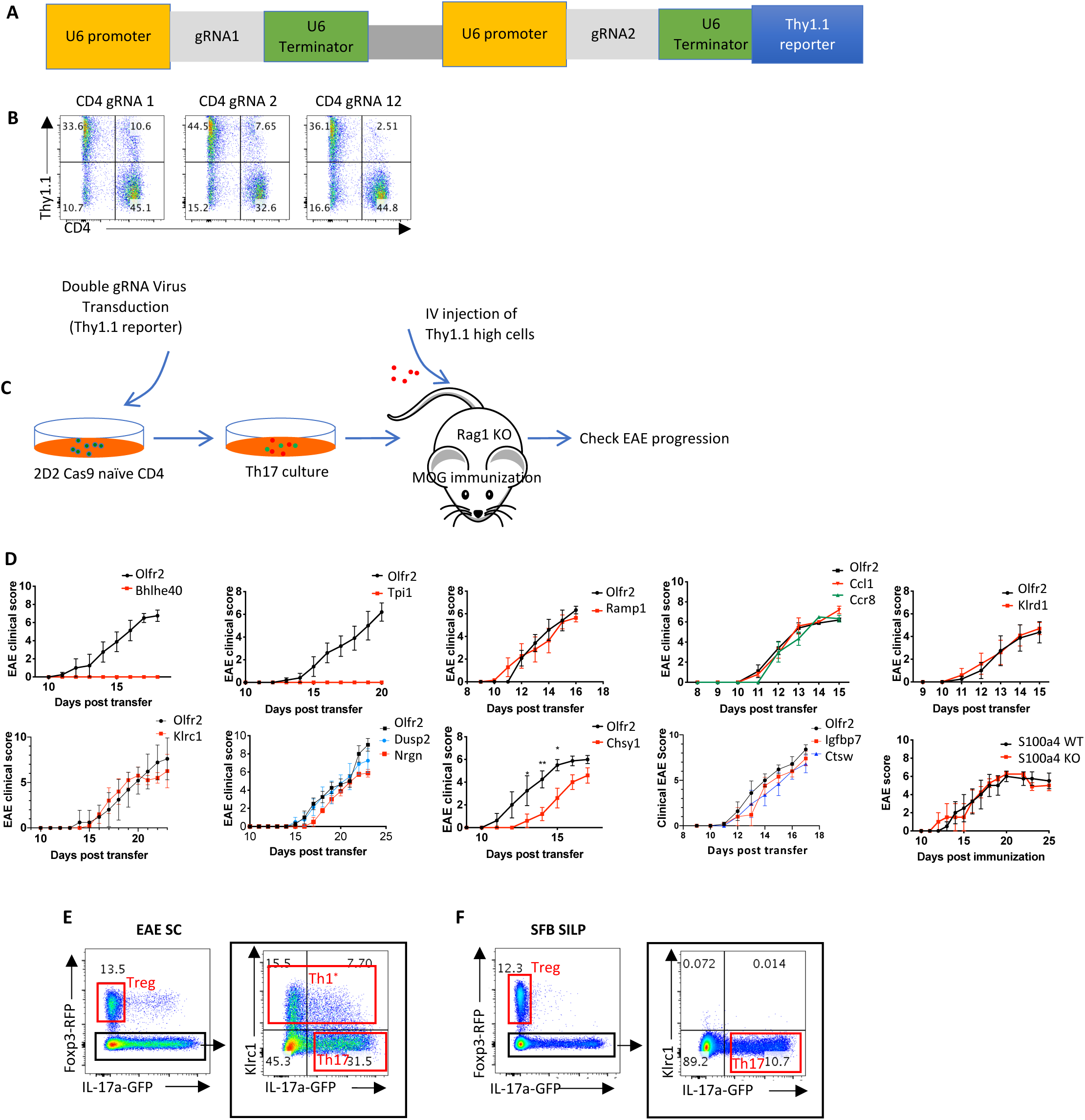
Tpi1 is essential for EAE-associated Th17 cell pathogenicity, related to Figure 1 and STAR Methods. (A) Schematic of the PSIN retroviral vector encoding the Thy1.1 reporter and two gRNAs. (B) Knockout efficiency with single compared to double gRNA vectors. In vitro cultured CD4^+^ T cells were infected with the retrovirus containing a single or two gRNAs targeting *Cd4,* and Cd4 expression was evaluated 72 h later. (C) Experimental design for functional screening of pathogenic genes in the transfer EAE model. Naïve 2D2 Cas9 CD4^+^ T cells were activated in vitro, infected with the double gRNA retrovirus, and differentiated into Th17 cells. Thy1.1 high cells were purified and transferred into *Rag1* KO mice (50k cells/mouse), which were immunized with MOG/CFA at the same time. EAE progression was monitored to evaluate the requirement for each gene in pathogenesis. (D) Clinical disease course of EAE in *Rag1* KO mice receiving control *(Olfr2* KO) 2D2 cells or candidate gene KO 2D2 cells, and in MOG-CFA-immunized *S100a4* KO and WT littermate control mice (6 pairs). The 2D2 transfer system was validated by failure of 2D2 cells deficient in *Bhlhe40*, a gene known to be essential for T cell pathogenicity, to induce EAE. At least 5 mice were used for each group, and each experiment was repeated at least twice. (E, F) Sorting strategy for bulk RNAseq analysis of EAE-associated Th17, Th1*, Treg (E) and SFB-induced Th17 and Treg cells (F).

**Figure S3.**
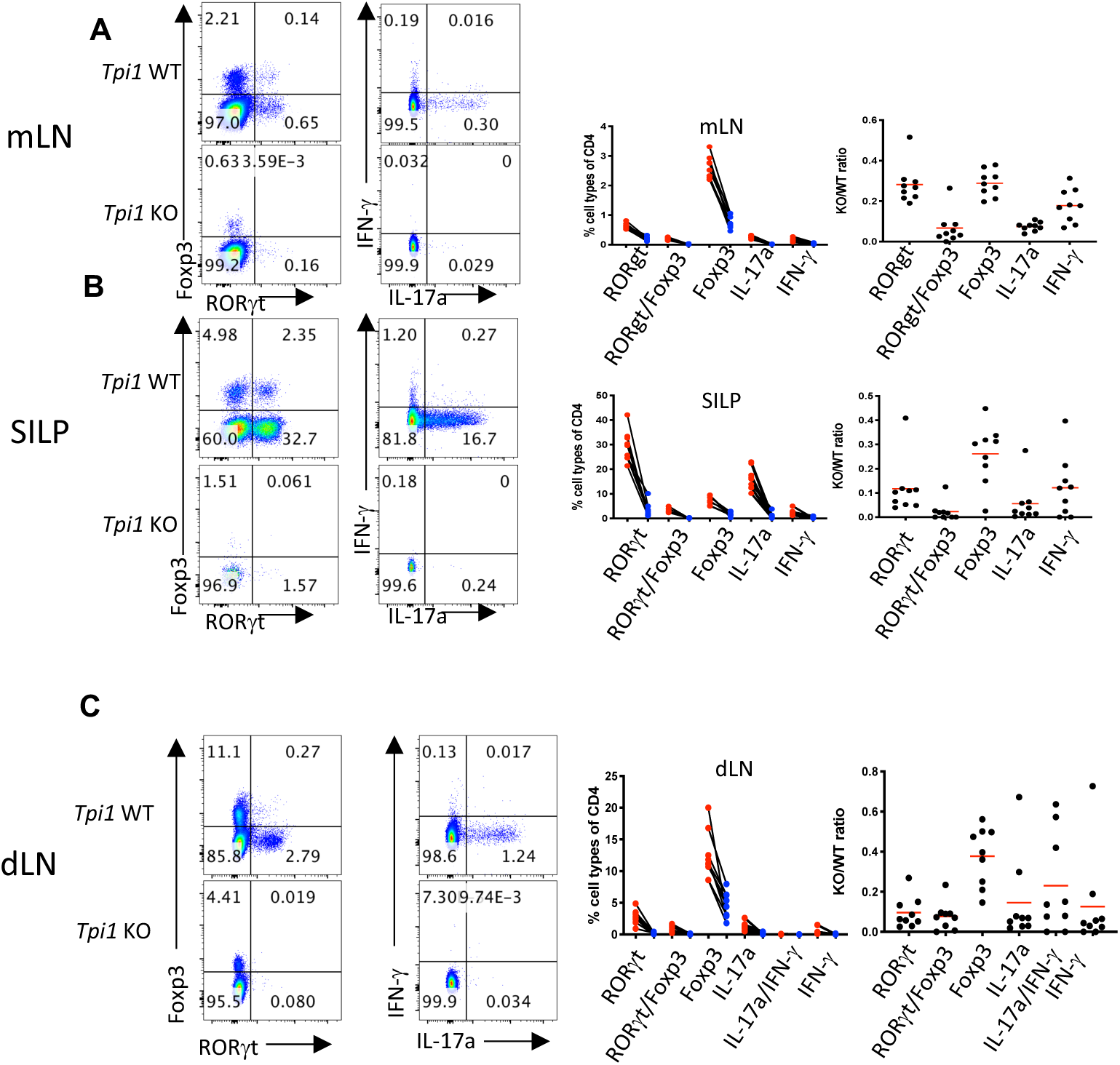
*Tpi1* is essential for the differentiation of all CD4^+^ T cells, related to Figure 1. (A, B) Impaired ROR*γ*t, Foxp3, IL-17a and IFN-*γ* expression by *Tpi1* mutant compared to WT CD4^+^ T cells in the mLN (A) and the SILP (B) of reconstituted mixed bone marrow chimeric mice colonized with SFB. Left panels, representative FACS plots showing ROR*γ*t/Foxp3 and IL-17a/IFN-*γ* expression in paired *Tpi1* WT (red) and KO (blue) CD4^+^ T cells in each mouse. Middle panels, frequencies of each CD4^+^ T cell subtype among the *Tpi1* WT and *Tpi1* KO cells. Right panels, *Tpi1* KO/*Tpi1* WT ratios for each CD4^+^ T cell subtype in mLN and SILP. (C) Impaired ROR*γ*t, Foxp3, IL-17a and IFN-*γ* expression in *Tpi1* mutant compared to WT CD4^+^ T cells in the dLN of mixed bone marrow reconstituted mice, at day 15 of MOG-induced EAE. The number of *Tpi1* KO CD4^+^ T cells obtained from SC was too low for a meaningful subtype analysis, and hence is not shown. Panels are organized as in (B).

**Figure S4.**
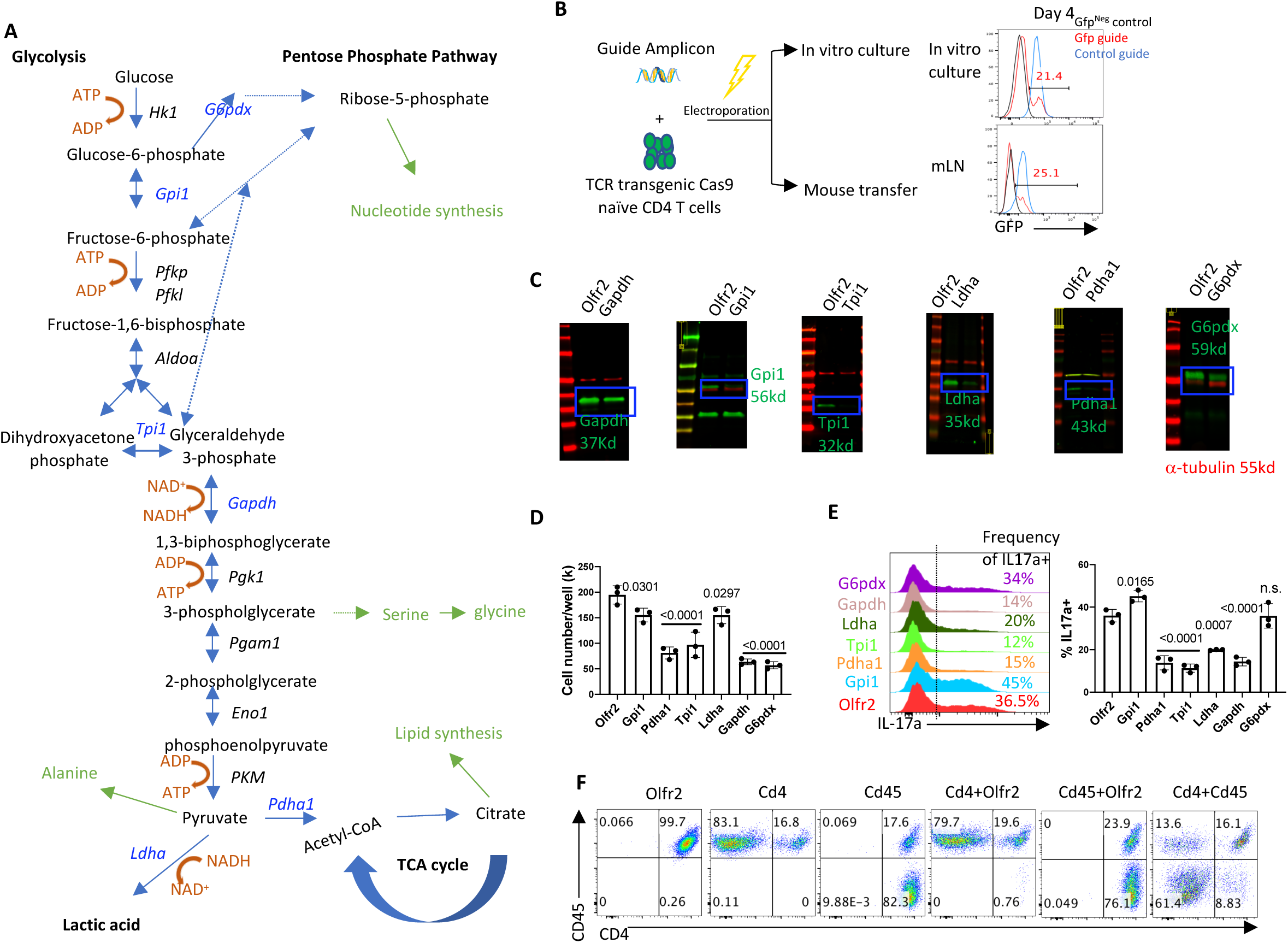
Electroporation-based CRISPR-mediated targeting of metabolic genes in naïve CD4^+^ T cells, related to Figure 2 and STAR Methods. (A) Map of metabolic pathways, showing relevant genes and metabolites in glycolysis, PPP and the TCA cycle. The biosynthetic processes from glycolysis metabolites are labeled in green. Genes that were targeted using CRISPR in this study are labeled in blue. Gpi1, Tpi1 and Ldha are the three enzymes that are expected to not be essential for glycolysis: (1) The reaction catalyzed by Gpi1 (glucose-6-phosphate to fructose-1,6-bisphosphate) can also be achieved by the PPP, making Gpi1 possibly redundant for this reaction; (2) Tpi1 controls the inter conversion of glyceraldehyde-3-phosphate and dihydroxyacetone phosphate, the two 3-carbon molecules generated by degrading fructose-1,6-phosphate. Loss of Tpi1 should lead to the loss of 50%, but not all of the glycolytic activity, and thus may be not lethal; (3) Ldha controls the conversion of pyruvate to lactic acid, and the regeneration of NAD^+^. However, pyruvate could still enter the TCA cycle in Ldha-deficient cells, for energy production and biomass synthesis. (B) Schematic of gene targeting in naïve T cells. Left, PCR amplicons encoding gRNAs were electroporated into Cas9-expressing naïve CD4^+^ T cells, which were subsequently cultured in vitro or transferred into recipient mice. Right, *in vivo* KO efficiency can be assessed by measuring the KO efficiency of the in vitro cultured Th17 cells. Cas9-expressing 7B8 naïve CD4^+^ T cells were electroporated with *Gfp* guide amplicons to target the Gfp that is co-expressed with Cas9 at the Rosa26 locus, and then cultured *in vitro* in Th17 conditions (IL6+TGF-*β*), or transferred into SFB-colonized mice. Gfp expression was examined 4 days later in the in vitro cultured Th17 cells (top) and the ex vivo cells isolated from the mLN (bottom). (C-E) Evaluation of the knockout efficiency of guides used in this study with in vitro Th17 cell culture. 8000 electroporated naïve CD4^+^ T cells were seeded in each well at the beginning of Th17 cell culture. Because gene function is related to cell growth and differentiation, we estimated the KO efficiency based on three variable factors at 120 h post electroporation: protein expression level (C), live cell number (D), and IL-17a production (E). Taking *Gapdh* targeting as an example, western blot data shows Gapdh protein expression decreased only ∼40%. However, the cell number was reduced ∼70%, and IL-17a production by live cells was further reduced more than 50% compared to the *Olfr2* control. Taken together, we estimate that the Gapdh guide achieved more than 80% knockout *in vitro*. Data are representative of three independent experiments. (F) Efficiency of gene inactivation following electroporation of naïve Cas9 CD4^+^ T cells with a mixture of two guide amplicons targeting *Cd4* and *Cd45* in cultured Th17 cells at 96 h. P values were determined using t tests.

**Figure S5.**
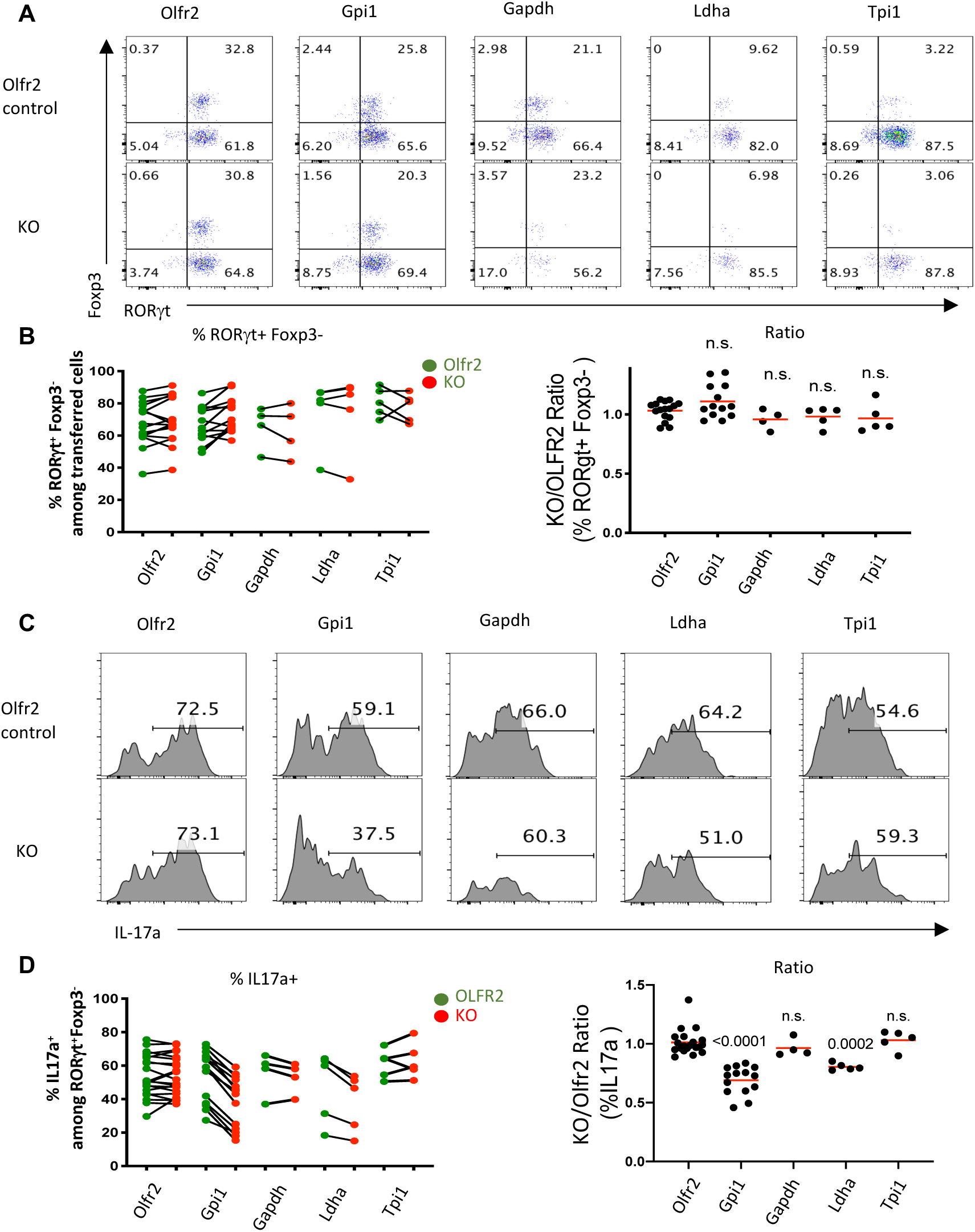
Differentiation of glycolysis gene-targeted 7B8 T cells in the SILP of SFB-colonized mice, related to Figure 2. (A) Representative FACS plots showing the expression of ROR*γ*t and Foxp3 in the *Olfr2* KO control cells and the co-transferred *Olfr2* control or glycolysis gene KO cells in the SILP at day 15 post transfer. (B) Left, summary of the Th17 frequencies (ROR*γ*t^+^ Foxp3^-^) in the *Olfr2* KO control cells (green) and the co-transferred *Olfr2* control or glycolysis gene KO cells (red), as in panel (A). Right, the KO/Olfr2 ratios of Th17 frequencies. (C) Representative FACS plots showing IL-17a expression in *Olfr2* control and co-transferred *Olfr2* control or glycolysis gene KO Th17 cells (ROR*γ*t^+^ Foxp3^-^ cells) isolated from the SILP at day 15 post transfer and following *in vitro* PMA/Ionomycin re-stimulation. (D) Left, summary of IL-17a expression based on results in panel (C). Right, KO/*Olfr2* ratios of IL-17a expression. There was consistent reduction of IL-17a in *Gpi1- or Ldha*-targeted, but not in *Olfr2*-targeted 7B8 cells isolated from the SILP and re-stimulated in vitro. While the reduced cytokine production observed in the *in vitro* restimulation assay supports the maintainence of Gpi1-deficient cells in the SILP, it is not necessarily contradictory to our previous claim that Gpi1-deficient 7B8 cells produce normal amounts of IL-17a (based on Il17a*^GFP/+^* assay (Figure 2H)), as *in vitro* restimulation was conducted in an artificial microenvironment with strong stimuli, wheareas *Il17a^GFP/+^* reporter cells report the true cytokine production in real physiological conditions. P values were determined using t tests.

**Figure S6.**
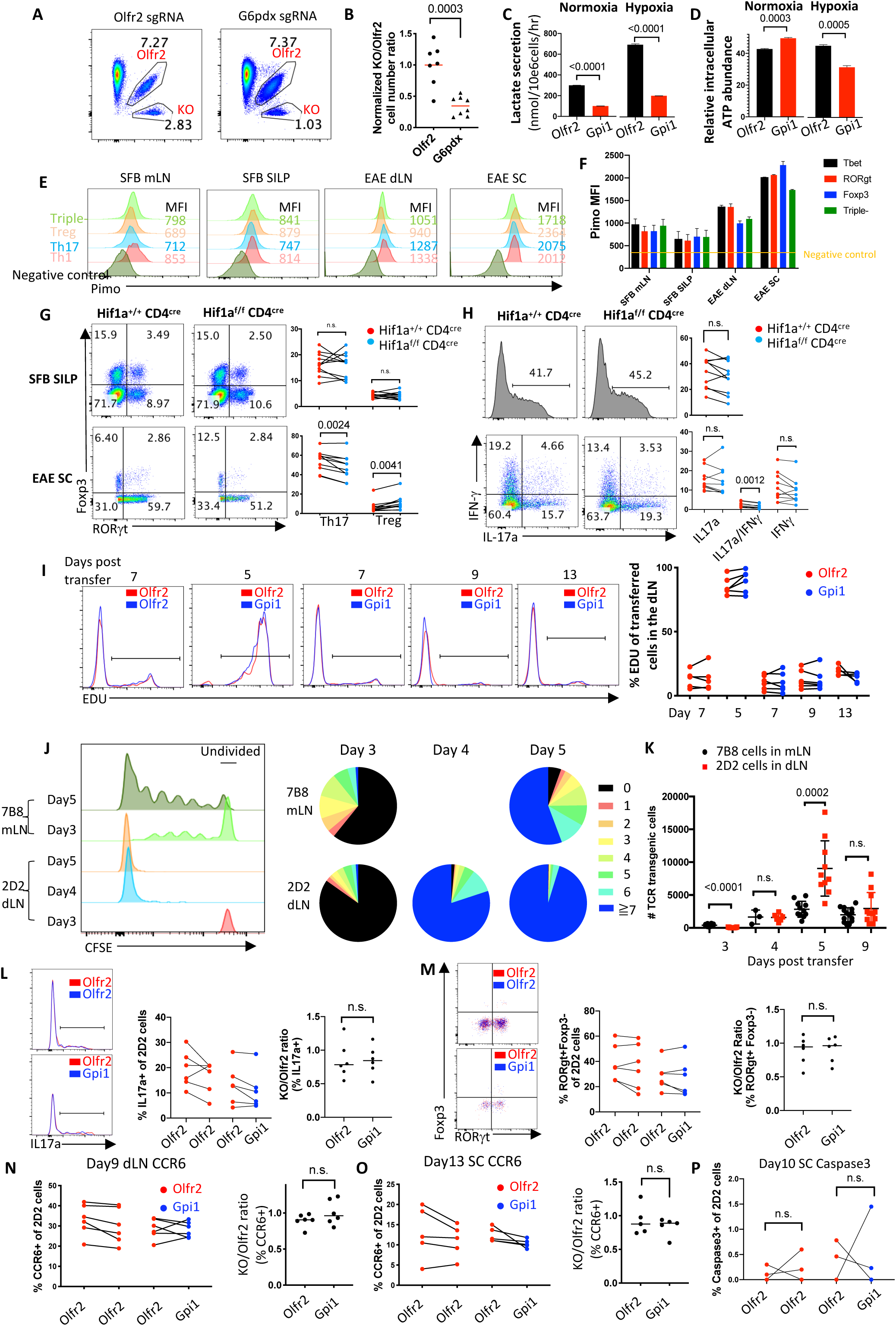
Hypoxic microenvironment in the EAE spinal cord impairs proliferation of *Gpi1*-deficient Th17 cells, related to Figure 6. (A, B) The PPP contributes to Th17 cell accumulation in the SC in EAE. (A) Representative FACS plots showing the frequencies of co-transferred *Olfr2* KO and *Olfr2* control or *G6pdx* KO 2D2 T cells in the EAE SC at day 15. (B) Compilation of KO/*Olfr2* cell number ratios as shown in panel A. (C) Lactate secretion of *in vitro* cultured *Olfr2* and *Gpi1* KO npTh17 cells in normoxia and hypoxia. Cells were cultured as in Figure 3A in normoxia. At 96 h cells were re-plated in new wells with fresh npTh17 medium in normoxia and hypoxia, and supernatants were collected 12 h later for lactate quantification by GC-MS. (D) Relative intracellular ATP abundance of in vitro cultured Olfr2 and Gpi1 KO npTh17 cells in normoxia and hypoxia. Cells were cultured as in Figure 3A in normoxic condition. At 96 h cells were re-plated in new wells with fresh npTh17 medium in normoxic and hypoxic condition, and ATP level was measured 12 h later. (E, F) Pimonidazole staining of different CD4^+^ T cell subsets in the SFB and EAE models. (E) Representative FACS plots showing pimonidazole staining of each subtype of CD4^+^ T cell in the mLN and the SILP of SFB-colonized healthy mice and in the dLN and the SC of EAE mice. Treg (Foxp3^+^ ROR*γ*t^-^ Tbet^-^), Th17 (ROR*γ*t^+^ Foxp3^-^Tbet^-^), Th1 (Tbet^+^ ROR*γ*t^-^ Foxp3^-^), Triple^-^ (ROR*γ*t^-^ Tbet^-^ Foxp3^-^). (F) Summarized pimonidazole staining based on panel C. (G) Transcription factor staining in CD4^+^ T cells from *Hif1a* WT and KO mice colonized with SFB (top, cells from SILP)) or subjected to EAE (bottom, from SC at day15). Left, representative FACS plots showing the frequencies of Th17 cells (ROR*γ*t^+^ Foxp3^-^) and Treg cells (Foxp3^+^ ROR*γ*t^-^) among total *Hif1a* WT and KO CD4^+^ T cells; Right, summary of transcription factor staining from two independent experiments. (H) Cytokine production in CD4^+^ T cells from *Hif1a* WT and KO CD4 T cells in the SFB (top) and EAE (bottom) models after in vitro PMA/ionomycin restimulation. Left, representative plots showing the frequencies of IL-17a producing cells among ROR*γ*t^+^ Foxp3^-^ *Hif1a* WT or KO cells in the SILP of SFB-colonized mice (top) and frequencies of IL-17a, IL-17a/IFN-*γ*, and IFN-*γ* producing cells among Foxp3^-^ *Hif1a* WT or KO cells in the SC at day 15 after EAE induction (bottom); Right, summary of cytokine production from two independent experiments.. (I) Left, representative plots showing EDU incorporation in co-transferred *Olfr2* and *Olfr2* control or *Gpi1* KO 2D2 cells at different days post transfer in the dLNs. Right, compiled data of one representative experiment. Two independent experiments were performed with the same conclusion. (J, K) Proliferation of dLN 2D2 cells in the EAE model and mLN 7B8 cells in SFB-colonized mice. Naïve 7B8 or 2D2 CD4^+^ T cells were labeled with CFSE and 100K cells were transferred into recipient mice that had been previously colonized with SFB or were immunized with MOG/CFA immediately after cell transfer. The mLN of the 7B8 model and dLN of the EAE model were collected at different time points for cell division analysis. (J) Left, representative FACS plot showing CFSE labeling of the 7B8 and 2D2 cells at different time points. Right, pie charts showing the frequencies of cells with different numbers of divisions. Division number was labeled in different colors. (K) Total cell number count of mLN 7B8 cells and dLN 2D2 cells at different time points. For each time point, 4-10 mice were used for each group. Two independent experiments were performed with the same conclusion. (L, M) IL-17a production (L) and ROR*γ*t/Foxp3 expression (M) in 2D2 cells from dLN at day 10 in the EAE model. Left, representative FACS plots showing the overlay of IL-17a expression (L) or ROR*γ*t/Foxp3 expression (M) in *Olfr2* control and co-transferred *Olfr2* KO or *Gpi1* KO 2D2 cells. Middle, compilation of frequencies of IL-17a^+^ (L) or ROR*γ*t^+^Foxp3^-^ cells (M) among *Olfr2* control and co-transferred *Olfr2* KO or *Gpi1* KO 2D2 cells. Right, the KO/Olfr2 ratios of the IL-17a^+^ frequencies (L) and ROR*γ*t^+^Foxp3^-^ frequencies (M). (N, O) Left, compiled data showing the frequencies of Ccr6^+^ cells among co-transferred *Olfr2* and *Olfr2* control or *Gpi1* KO 2D2 cells in the dLN at day 9 (N) or in the SC at day 13 (O) post transfer and EAE induction. Right, the KO/Olfr2 ratios of the Ccr6^+^ frequencies. Two independent experiments were performed with the same conclusion. (P) Compiled data showing the frequencies of active Caspase3^+^ cells among co-transferred *Olfr2* and *Olfr2* control or *Gpi1* KO 2D2 cells in the SC at day 10 post transfer and EAE induction. n=10. P values of panel B, C, D, K, L, M, N, and O were determined using t tests. P values of panel G, H, and P, were determined using paired t tests.

## STAR Methods

### RESOURCE AVAILABILITY

### LEAD CONTACT AND MATERIALS AVAILABILITY

Further information and requests for resources and reagents should be directed to and will be fulfilled by the Lead Contact, Dan R. Littman.

#### Materials Availability

This study did not generate new unique reagents

#### Data and Code Availability

The bulk RNAseq data and the scRNA seq data supporting the current study have been deposited in GEO (accession number GSE141006).

### EXPERIMENTAL MODEL AND SUBJECT DETAILS

#### Mouse Strains

C57BL/6 mice were obtained from Taconic Farms or the Jackson Laboratory. All transgenic animals were bred and maintained in the animal facility of the Skirball Institute (NYU School of Medicine) in specific-pathogen-free (SPF) conditions. *Il-23r^GFP^* mice (Awasthi et al., 2009) were provided by M. Oukka. *S100a4* knockout mice (Li et al., 2010) were provided by Dr. Anne R. Bresnick. *Cas9^Tg^* mice (Platt et al., 2014) were obtained from Jackson Laboratory (JAX; B6J.129(Cg)-Gt(ROSA)26Sortm1.1(CAG-cas9*,-EGFP)Fezh/J), and maintained on the Cd45.1 background (JAX; B6. SJL-Ptprca Pepcb/BoyJ). SFB-specific TCRTg (7B8) mice (Yang et al., 2014), MOG-specific TCRTg (2D2) mice (Bettelli et al., 2003), and *Helicobacter hepaticus*-specific TCRTg (HH7-2) mice (Xu et al., 2018) were previously described. They were crossed to the Cas9^Tg^ Cd45.1/1 mice to generate 7B8 *Cas9 ^Tg^*, 2D2 *Cas9 ^Tg^*, and HH7-2 *Cas9 ^Tg^* mice with either Cd45.1/1 or Cd45.1/2 congenic markers. 7B8 Cas9 CD45.1/2 mice were further crossed with *IL17a^GFP/GFP^* reporter mice (JAX; C57BL/6-Il17atm1Bcgen/J) to generate 7B8 *Cas9 IL17a^GFP/^*^+^ Cd45.1/2 or Cd45.2/2 mice. Female *Foxp3^RFP/RFP^* reporter mice (Jax; C57BL/6-Foxp3^tm1Flv^/J) were crossed with male *IL17a^GFP/GFP^* reporter mice to generate *IL17a^GFP/+^ Foxp3^RFP^* male mice for bulk RNAseq. *Hif1a^f/f^* mice (Jax; B6.129-Hif1a^tm3Rsjo^/J) were crossed with *CD4^cre^* mice (Jax; B6.Cg-Tg(Cd4-cre)1Cwi/BfluJ) and CD45.1/1 mice to generate *Hif1a^+/+^ CD4^cre^* CD45.1/1 and *Hif1a^f/f^ CD4^cre^* CD45.2/2 littermates. *Tpi1^f/+^* mice generated by crossing *Tpi1^tm1a(EUCOMM)Wtsi^* mice (Wellcome Trust Sanger Institute; B6Brd;B6N-Tyr^c-Brd^ Tpi1^tm1a(EUCOMM)Wtsi^/WtsiCnbc) with a FLP deleter strain (Jax; B6.129S4-Gt(ROSA)26Sor^tm1(FLP1)Dym^/RainJ) were further crossed with CD4^cre^ and CD45.1/1 strains to generate *Tpi1^+/+^Cd4^cre^* and *Tpi1^f/f^CD4^cre^* mice with different congenic markers.

### METHOD DETAILS

#### Flow cytometry

For transcription factor and cytokine staining of *ex vivo* cells, single cell suspensions were incubated for 3h in RPMI with 10% FBS, phorbol 12-myristate 13-acetate (PMA) (50 ng/ml; Sigma), ionomycin (500 ng/ml;Sigma) and GolgiStop (BD). Cells were then resuspended with surface-staining antibodies in HEPES-buffered HBSS. Staining was performed for 20-30 min on ice. Surface-stained cells were washed and resuspended in live/dead fixable blue (ThermoFisher) for 5 min prior to fixation. Cells were treated using the FoxP3 staining buffer set from eBioscience according to the manufacturer’s protocol. Intracellular stains were prepared in 1X eBioscience permwash buffer containing anti-CD16/anti-CD32, normal mouse IgG, and normal rat IgG Staining was performed for 30-60 min on ice.

For cytokine analysis of *in vitro* cultured Th17 cells, cells were incubated in the restimulation buffer for 3 h. After surface and live/dead staining, cells were treated using the Cytofix/Cytoperm buffer set from BD Biosciences according to the manufacturer’s protocol. Intracellular stains were prepared in BD permwash in the same manner used for transcription factor staining. Flow cytometric analysis was performed on an LSR II (BD Biosciences) or an Aria II (BD Biosciences) and analyzed using FlowJo software (Tree Star).

#### Retroviral transduction and 2D2 TCRtg Cas9^Tg^ T cell-mediated EAE

The protocol described in this section was only used for experiments shown in Figure 2F and Figure S2A-D. sgRNAs were designed using the Crispr guide design software (http://crispr.mit.edu/). sgRNA encoding sequences were cloned into the retroviral sgRNA expression vector pSIN-U6-sgRNAEF1as-Thy1.1-P2A-Neo that has been reported before (Ng et al., 2019). To make the double sgRNA vector for experiments shown in Figures S2A-D, the sgRNA transcription cassette, ranging from the U6 promoter to the U6 terminator, was amplified (primer sequences can be found in the Key Resources Table) and cloned into the pSIN vector after the first sgRNA encoding cassette. For the EAE experiment shown in Figure 2F, only one sgRNA was used for each virus. Retroviruses were packaged in PlatE cells by transient transfection using TransIT 293 (Mirus Bio).

Tissue culture plates were coated with polyclonal goat anti-hamster IgG (MP Biomedicals) at 37 °C for at least 3 h and washed 3× with PBS before cell plating. FACS-sorted CD4^+^CD8^−^CD25^−^CD62L^+^CD44^low^ naïve 2D2 TCRtg *Cas9^Tg^* T cells were seeded for 24 h in T cell medium (RPMI supplemented with 10% FCS, 2mM *β*-mercaptoethanol, 2mM glutamine), along with anti-CD3 (BioXcell, clone 145-2C11, 0.25µg/ml) and anti-CD28 (BioXcell, clone 37.5.1, 1µg/ml) antibodies. Cells were then transduced with the pSIN virus by spin infection at 1200 × g at 30 °C for 90 min in the presence of 10µg/ml polybrene (Sigma). Viral supernatants were removed 12 h later and replaced with fresh medium containing anti-CD3/anti-CD28, 20ng/ml IL-6 and 0.3ng/ml TGF-*β*. Cells were lifted off 48 h later and supplemented with IL-2 for another 48hours. Thy1.1^high^ cells were then sorted with the ARIA and i.v. injected into *Rag1*^-/-^ mice (50k cells/mouse). Mice were then immunized subcutaneously on day 0 with 100µg of MOG35-55 peptide, emulsified in CFA (Complete Freund’s Adjuvant supplemented with 2mg/mL *Mycobacterium tuberculosis* H37Ra), and injected i.p. with 200 ng pertussis toxin (Calbiochem). The EAE scoring system was as follows: 0-no disease, 1-Partially limp tail; 2-Paralyzed tail; 3-Hind limb paresis, uncoordinated movement; 4-One hind limb paralyzed; 5-Both hind limbs paralyzed; 6-Hind limbs paralyzed, weakness in forelimbs; 7-Hind limbs paralyzed, one forelimb paralyzed; 8-Hind limbs paralyzed, both forelimbs paralyzed; 9-Moribund; 10-Death.

#### sgRNA guide amplicon electroporation of naïve CD4^+^ T cells

sgRNA-encoding sequences were cloned into the pSIN vector (single sgRNA in one vector). The sgRNA encoding cassette, ranging from the U6 promoter to the U6 terminator, was amplified with 34 cycles of PCR with Phusion (NEB), and purified with Wizard SV Gel and PCR Clean-up System (Promega), and concentrated with a standard ethanol DNA precipitation protocol. Amplicons were resuspended in a small volume of H2O and kept at −80 °C until use. Amplicon concentration was measured with Qubit dsDNA HS assay kit (Thermofisher)

The Amaxa P3 Primary Cell 4D-Nucleofector X kit was used for T cell electroporation. 2 million FACS-sorted naïve CD4^+^ *Cas9^Tg^* cells were pelleted and resuspended in 100ul P3 primary cell solution supplemented with 0.56ug sgRNA amplicon (for single gene knockout), or equal amounts of sgRNA amplicons (0.56ug each for targeting two genes simultaneously in Figures 3D, and 4E). For GFP KO (Figure S4B), a mixture of three pSIN vectors at equal amount containing different sgRNA-encoding sequences (all targeting GFP) was used as template for PCR amplifaction, and 0.56ug of amplicon was used for electroporation. Cell suspensions were transferred into a Nucleocuvette Vessel for electroporation in the Nucleofector X unit, using program “T cell. Mouse. Unstim.”. Cells were then rested in 500 ul mouse T cell nucleofector medium (Lonza) for 30 min before in vitro culture or transfer into mice.

In vitro cell culture – 96 well flat-bottom tissue culture plates (Corning) were coated with polyclonal goat anti-hamster IgG (MP Biomedicals) at 37 °C for at least 3 h and washed 3× with PBS before cell plating. Electroporated cells were plated into each well containing 200ul T cell culture medium (glucose-free RPMI (ThermoFisher) supplemented with 10% FCS (Atlanta Biologicals), 4g/L glucose, 4mM glutamine, 50uM *β*-mercaptoethanol, 0.25µg/ml anti-CD3 (BioXcell, clone 145-2C11) and 1µg/ml anti-CD28 (BioXcell, clone 37.5.1) antibodies). For npTh17 culture, 20ng/ml IL-6 and 0.3ng/ml TGF-*β* were further added into the T cell medium. For pTh17 cell culture, 20ng/ml IL-6, 20ng/ml IL-23, and 20ng/ml IL-1*β* were further added into the T cell medium. Cells were then cultured for 96 h or 120 h in normoxia or hypoxia (3% oxygen) before metabolic experiments or cell number counting/western/FACS profiling, respectively.

T cell transfer into mice – TCR transgenic *Cas9^Tg^* naïve CD4^+^ T cells (7B8 TCRtg for SFB model, 2D2 TCRtg for EAE model, and HH7-2 TCRtg for colitis model) with different isotype markers were electroporated at the same time. Equal numbers of isotype-distinct electroporated resting cells were mixed, pelleted and resuspended in dPBS supplemented with 1% filtered FBS before retro-orbital injection into recipient mice (100,000 cells in total for each mouse). The transferred cells were isolated from SILP of the SFB model, SC of the EAE model, and the LILP of the colitis model at day 15 post transfer, unless otherwise indicated.

#### SFB model

C57BL/6 mice were purchased from Jackson Laboratory and maintained in SFB-free SPF cages. Fresh feces of SFB mono-colonized mice maintained in our gnotobiotic animal facility were collected and frozen at −80 °C until use. Fecal pellets were meshed through a 100uM strainer (Corning) in cold sterile dPBS (Hyclone). 300 µl feces suspension, equal to the amount of 1/3 pellet, was administered to each mouse by oral gavage. Mice were inoculated one more time at 24 h. For the transfer experiment, electroporated cells were injected 96 h after the initial gavage.

#### Colitis model

G. *hepaticus* was provided by J. Fox (MIT) and grown on blood agar plates (TSA with 5% sheep blood, Thermo Fisher) as previously described (Xu et al., 2018). Briefly, *Rag1*^-/-^ mice were colonized with *H. hepaticus* by oral gavage on days 0 and 4 of the experiment. At day 7, electroporated naïve HH7-2 TCRtg *Cas9^Tg^* CD4^+^ T cells were adoptively transferred into the *H. hepaticus*-colonized recipients. In order to confirm *H. hepaticus* colonization, *H. hepaticus*-specific 16S primers were used on DNA extracted from fecal pellets. Universal 16S were quantified simultaneously to normalize *H. hepaticus* colonization of each sample.

#### EAE model

Right after the transfer of electroporated 2D2 *Cas9^Tg^* cells, mice were immunized subcutaneously on day 0 with 100µg of MOG35-55 peptide, emulsified in CFA (Complete Freund’s Adjuvant supplemented with 2mg/mL Mycobacterium tuberculosis H37Ra), and injected i.p. on days 0 and 2 with 200 ng pertussis toxin (Calbiochem).

#### Isolation of lymphocytes from lamina propria, spinal cord, and lymph nodes

For isolating mononuclear cells from the lamina propria, the intestine (small and/or large) was removed immediately after euthanasia, carefully stripped of mesenteric fat and Peyer’s patches/cecal patch, sliced longitudinally and vigorously washed in cold HEPES buffered (25mM), divalent cation-free HBSS to remove all fecal traces. The tissue was cut into 1-inch fragments and placed in a 50ml conical containing 10ml of HEPES buffered (25mM), divalent cation-free HBSS and 1 mM of fresh DTT. The conical was placed in a bacterial shaker set to 37 °C and 200rpm for 10 minutes. After 45 seconds of vigorously shaking the conical by hand, the tissue was moved to a fresh conical containing 10ml of HEPES buffered (25mM), divalent cation-free HBSS and 5 mM of EDTA. The conical was placed in a bacterial shaker set to 37 °C and 200rpm for 10 minutes. After 45 seconds of vigorously shaking the conical by hand, the EDTA wash was repeated once more in order to completely remove epithelial cells. The tissue was minced and digested in 5-7ml of 10% FBS-supplemented RPMI containing collagenase (1 mg/ml collagenaseD; Roche), DNase I (100 µg/ml; Sigma), dispase (0.05 U/ml; Worthington) and subjected to constant shaking at 155rpm, 37 °C for 35 min (small intestine) or 55 min (large intestine). Digested tissue was vigorously shaken by hand for 2 min before adding 2 volumes of media and subsequently passed through a 70 µm cell strainer. The tissue was spun down and resuspended in 40% buffered percoll solution, which was then aliquoted into a 15ml conical. An equal volume of 80% buffered percoll solution was underlaid to create a sharp interface. The tube was spun at 2200rpm for 22 min at 22 °C to enrich for live mononuclear cells. Lamina propria (LP) lymphocytes were collected from the interface and washed once prior to staining.

For isolating mononuclear cells from spinal cords during EAE, spinal cords were mechanically disrupted and dissociated in RPMI containing collagenase (1 mg/ml collagenaseD; Roche), DNase I (100 µg/ml; Sigma) and 10% FBS at 37 °C for 30 min. Leukocytes were collected at the interface of a 40%/80% Percoll gradient (GE Healthcare).

For isolating mononuclear cells from lymph nodes, mesenteric lymph nodes (for the SFB model), and draining lymph nodes (Inguinal, axillary, and brachial lymph nodes for the EAE model) were mechanically dissected and minced in RPMI containing 10% FBS and 25mM HEPES through 40uM strainer. Cells were collected for further analysis.

#### Generation of bone marrow chimeric reconstituted mice

Bone marrow mononuclear cells were isolated from donor mice by flushing the long bones. To generate *Tpi1^WT^/ Tpi1^KO^* chimeric reconstituted mice, *Tpi1^+/+^ Cd4^Cre^* CD45.1/2 and *Tpi1^f/f^ Cd4^Cre^* CD45.2/2 mice were used as donors. To generate *Hif1a^WT^/Hif1a^KO^* chimeric reconstituted mice, *Hif1a^+/+^ Cd4^Cre^* CD45.1/2 and *Hif1a^f/f^ Cd4^Cre^* CD45.1/1 mice were used as donors. To generate *Il23r^het^/Il23r^KO^* chimeric reconstituted mice, *Il23r^Gfp/+^* CD45.1/2 *and Il23r^Gfp/Gfp^* CD45.1/1 mice were used as donors. Red blood cells were lysed with ACK Lysing Buffer, and CD4/CD8 T cells were labeled with CD4/CD8 magnetic microbeads and depleted with a Miltenyi LD column. The remaining cells were resuspended in PBS for injection in at a 1:1 ratio (WT: KO).

4×10^6^ cells were injected intravenously into 6 week old mice (CD45.1/1 in Tpi1 experiment; Rag1^-/-^ in Hif1a experiment) that were irradiated 4h before reconstitution using 1000 rads/mouse (2×500rads, at an interval of 3h, at X-RAD 320 X-Ray Irradiator). Peripheral blood samples were collected and analyzed by FACS 8 weeks later to check for reconstitution, after which SFB colonization or EAE induction were performed.

#### Pimonidazole HCl labeling

Pimonidazole HCl (hypoxyprobe) was retro orbitally injected into mice at the dose of 100mg/kg under general anesthesia (Ketamine 100mg/Kg, Xylazine 15mg/Kg). 3 h later, tissues were dissected and processed to get the mononuclear cell fraction. After surface/live/dead staining, cells were treated using the FoxP3 staining buffer set from eBioscience according to the manufacturer’s protocol. Intracellular staining with anti-ROR*γ*t, Foxp3, Tbet, and biotin-pimonidazole adduct antibodies were performed for 60min on ice in 1X eBioscience permwash buffer containing normal mouse IgG (50μg/ml), and normal rat IgG (50μg/ml). The pimonidazole labeling was visualized with streptavidin APC staining in the same buffer.

#### EdU incorporation

Two doses of EdU (50mg/kg for each) were i.p. injected into MOG/CFA-immunized mice in 12 h intervals at days 4, 6, 8, 9, and 12 post immunization. Draining lymph nodes and/or spinal cords were dissected 12 h post the second injection. Detection of EdU incorporation into the DNA was performed with EdU-click 647 Kit (Baseclick) according to the manufacturer’s instructions. In brief, surface-stained cells were fixed in 100μl fixative solution, and permeabilized in 100μl permeabilization buffer. Cells were pelleted and incubated for 30min in 220μl EdU reaction mixture containing 20μl permeabilization buffer, 20μl buffer additive, 1μl dye azide, 4μl catalyst solution, and 175μl dPBS, before 3x wash and FACS analysis.

#### CFSE labeling

Sorted naïve 7B8 or 2D2 CD45.1/1 CD4 T cells were stained with CFSE cell proliferation kit (Life Technology). Labeled cells were administered into each congenic CD45.2/2 recipient mouse by retro-orbital injection. Mice receiving 7B8 cells were gavaged with SFB mono-feces twice on days −4 and −3. Mice receiving 2D2 cells were immunized for EAE induction immediately after cell transfer. mLNs of the SFB-colonized mice were collected at 72h and 120h, and dLN of 2D2-injected mice were collected at 72h, 96h and 120h post transfer for cell division analysis.

#### Bulk RNA sequencing

Total RNA from sorted target cell populations was isolated using TRIzol LS (Invitrogen) followed by DNase I (Qiagen) treatment and cleanup with RNeasy Plus Micro kit (Qiagen; 74034). RNA quality was assessed using Pico Bioanalyser chips (Agilent). RNASeq libraries were prepared using the Clontech Smart-Seq Ultra low RNA kit (Takara Bio) for cDNA preparation, starting with 550 pg of total RNA, and 13 cycles of PCR for cDNA amplification. The SMARTer Thruplex DNA-Seq kit (Takara Bio) was used for library prep, with 7 cycles of PCR amplification, following the manufacturer’s protocol. The amplified library was purified using AMPure beads (Beckman Coulter), quantified by Qubit and QPCR, and visualized in an Agilent Bioanalyzer. The libraries were pooled equimolarly, and run on a HiSeq 4000, as paired end reads, 50 nucleotides in length.

#### Single cell RNAseq

Libraries from isolated single cells were generated based on the Smart-seq2 protocol (Picelli et al., 2014) with the following modifications. In brief, RNA from sorted single cells was used as template for oligo-dT primed reverse transcription with Superscript II Reverse Transcriptase (Thermo Fisher), and the cDNA library was generated by 20 cycle PCR amplification with IS PCR primer using KAPA HiFi HotStart ReadyMix (KAPA Biosystems) followed by Agencourt AMPure XP bead purification (Beckman Coulter) as described. Libraries were tagmented using the Nextera XT Library Prep kit (Illumina) with custom barcode adapters (sequences available upon request). Libraries with unique barcodes were combined and sequenced on a HiSeq2500 in RapidRun mode, with either 2×50 or 1×50 nt reads.

#### Stable-isotope tracing, metabolite secretion and kinetic flux profiling by GC-MS

*In vitro* cultured np/pTh17 cells were collected at 96h, washed twice in glucose-free RPMI (ThermoFisher, 11879020) containing 10% dialyzed FBS (ThermoFisher), and re-plated in anti-hamster IgG-coated 96-well plates at 60,000/well in np/pTh17 medium (glucose-free RPMI containing 10% dialyzed FBS, 2mM *b*-mercaptoethanol, 4mM glutamine, anti-CD3, anti-CD28, and 4g/L isotype labeled glucose). For KA-treated samples, *Olfr2* KO cells were previously treated with 30 µM KA before collection and re-plating. The re-plated cells were cultured in Th17 medium supplemented with 30 µM KA.

To measure relative PPP activity, ^13^C1,2-glucose (Cambridge Isotope Laboratories) was added into the Th17 medium. Re-plated cells were further cultured 12 h before metabolite extraction and GC-MS analysis. After a brief wash with 0.9% ice-cold saline solution to remove media contamination, cellular metabolites were extracted using a pre-chilled methanol/water/chloroform extraction method, as previously described (Metallo et al., 2011). Aqueous and inorganic layers were separated by cold centrifugation for 15 min. The aqueous layer was transferred to sample vials (Agilent 5190-2243) and evaporated to dryness using a SpeedVac (Savant Thermo SPD111V).

For lactate secretion analysis, re-plated cells were cultured in normoxia or hypoxia for 12 h. media was collected and centrifuged at 1,000 x g to pellet cellular debris. 5 µL of supernatant was extracted in 80% ice-cold methanol mixture containing isotope-labeled internal standards and evaporated to dryness.

Metabolite abundance [Xm] was determined using the following equation:

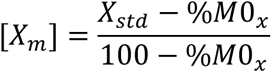

where Xstd is the molar amount of isotope-labeled standard (e.g. 5 nmol lactate) and %M0x is the relative abundance of unlabeled metabolite X corrected for natural isotope abundance.

To quantify the metabolite secretion flux (Fluxm), the molar change in extracellular metabolites was determined by calculating the difference between conditioned and fresh media and normalizing to changes in cell density:

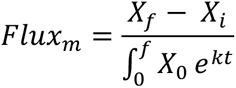

where Xf and Xi are the final and initial molar amounts of metabolite, respectively, X0 is the initial cell density, k is the exponential growth rate (hr^-1^), and t is the media conditioning time.

For kinetic flux profiling, npTh17 medium was supplemented with 4g/L ^13^C6-glucose (Cambridge Isotope Laboratories). Cells were collected at various time points. Unlabeled metabolite (M0) versus time (t) were plotted and the data were fitted to an equation as previously described (Yuan et al., 2008) to quantify the metabolite turnover rate (t^-1^):

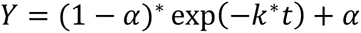

where *α*is the percentage of unlabeled (M0) metabolite at steady-state, k is the metabolite turnover rate and t is time.

Metabolite flux was quantified by multiplying the decay rate by the intracellular metabolite abundance.

Metabolite derivatization using MOX-tBDMS was conducted as previously described (Lewis et al., 2014). Derivatized samples were analyzed by GC-MS using a DB-35MS column (30m x 0.25mm i.d. x 0.25 µm) installed in an Agilent 7890B gas chromatograph (GC) interfaced with an Agilent 5977B mass spectrometer as previously described (Grassian et al., 2014) and corrected for natural abundance using in-house algorithms adapted from (Fernandez et al., 1996).

#### Intracellular ATP measurement

npTh17 cells were cultured in vitro in normoxic condition for 96 h before being collected and washed twice in glucose-free RPMI (ThermoFisher) containing 10% FBS (Atlanta Biologicals), and re-plated in anti-hamster IgG-coated 96-well plates at 20,000/well in 200μl npTh17 medium (glucose-free RPMI containing 10% dialyzed FBS, 2mM *β*-mercaptoethanol, 4mM glutamine, anti-CD3, anti-CD28, and 4g/L glucose). 12 hours later, cell numbers were determined in replicate wells, and ATP measurements were made using CellTiter-Glo® Luminescent Cell Viability Assay kit (Promega) according to instructions.

#### LC-MS analysis

For glucose consumption analysis, supernatants of npTh17 cells, or cell free control, were collected at 24h post cell re-plating. Metabolites were extracted for LC-MS analysis to measure glucose and lactate quantities. Glucose consumption rate and lactate production rate was quantified on a relative basis by calculating the ion count difference between conditioned and fresh media and normalizing to changes in cell density.

Samples were subjected to an LC-MS analysis to detect and quantify known peaks. A metabolite extraction was carried out on each sample with a previously described method (Pacold et al., 2016). In brief, samples were extracted by mixing 10 µL of sample with 490 µL of extraction buffer (81.63% methanol (Fisher Scientific) and 500 nM metabolomics amino acid mix standard (Cambridge Isotope Laboratories, Inc.)) in 2.0mL screw cap vials containing ∼100 µL of disruption beads (Research Products International, Mount Prospect, IL). Each was homogenized for 10 cycles on a bead blaster homogenizer (Benchmark Scientific, Edison, NJ). Cycling consisted of a 30 sec homogenization time at 6 m/s followed by a 30 sec pause. Samples were subsequently spun at 21,000 g for 3 min at 4 °C. A set volume of each (360 µL) was transferred to a 1.5 mL tube and dried down by speedvac (Thermo Fisher, Waltham, MA). Samples were reconstituted in 40 µL of Optima LC/MS grade water (Fisher Scientific, Waltham, MA).

Samples were sonicated for 2 mins, then spun at 21,000g for 3min at 4°C. Twenty microliters were transferred to LC vials containing glass inserts for analysis.

The LC column was a MilliporeTM ZIC-pHILIC (2.1 x150 mm, 5 µm) coupled to a Dionex Ultimate 3000TM system and the column oven temperature was set to 25 °C for the gradient elution. A flow rate of 100 µL/min was used with the following buffers; A) 10 mM ammonium carbonate in water, pH 9.0, and B) neat acetonitrile. The gradient profile was as follows; 80-20%B (0-30 min), 20-80%B (30-31 min), 80-80%B (31-42 min). Injection volume was set to 1 µL for all analyses (42 min total run time per injection).

MS analyses were carried out by coupling the LC system to a Thermo Q Exactive HFTM mass spectrometer operating in heated electrospray ionization mode (HESI). Method duration was 30 min with a polarity switching data-dependent Top 5 method for both positive and negative modes. Spray voltage for both positive and negative modes was 3.5kV and capillary temperature was set to 320°C with a sheath gas rate of 35, aux gas of 10, and max spray current of 100 µA. The full MS scan for both polarities utilized 120,000 resolution with an AGC target of 3e6 and a maximum IT of 100 ms, and the scan range was from 67-1000 m/z. Tandem MS spectra for both positive and negative mode used a resolution of 15,000, AGC target of 1e5, maximum IT of 50 ms, isolation window of 0.4 m/z, isolation offset of 0.1 m/z, fixed first mass of 50 m/z, and 3-way multiplexed normalized collision energies (nCE) of 10, 35, 80. The minimum AGC target was 1e4 with an intensity threshold of 2e5. All data were acquired in profile mode.

#### Seahorse Analysis

OCR and ECAR measurements were performed using a Seahorse XFe96 analyzer (Seahorse Biosciences). Seahorse XFe96 plates were coated with 0.56ug Cell-Tak (Corning) in 24µl coating buffer (8µl H2O + 16µl 0.1M NaHCO3 (PH 8.0)) overnight at 4 °C, and washed 3x with H2O before use. *In vitro* cultured Th17 cells were collected at 96 h, washed twice in Seahorse RPMI media PH7.4 (Agilent, 103576-100) and seeded onto the coated plates at 100,000 per well. To measure the OCR and ECAR of KA-treated cells, 30μM KA (Santa Cruz) was added into *Olfr2* control Th17 cell samples at the beginning of the Seahorse measurement. Respiratory rates were measured in RPMI Seahorse media, pH7.4 (Agilent) supplemented with 10 mM glucose (Agilent) and 2 mM glutamine (Agilent) in response to sequential injections of oligomycin (1 µM), FCCP (0.5 µM) and antimycin/rotenone (1 µM) (all Cayman Chemicals, Ann Arbor, MI, USA).

#### Western blotting

Whole cell extracts were fractionated by SDS-PAGE and transferred to nitrocellulose membrane by wet transfer for 4 h on ice at 60 V. Blots were blocked in TBS blocking buffer (Licor) and then stained overnight with primary antibody at 4°C. The next day, membranes were washed three times in TBST (TBS and 0.1% Tween-20), stained with fluorescently conjugated secondary antibodies (Licor) at 1:10,000 dilution in TBS blocking buffer for 1 h at room temperature, and then imaged in the 680- and 800-nm channels. Antibodies used: Gapdh (CST), Gpi1 (Thermo Fisher), Ldha (CST), Pdha1 (Abcam), Tpi1 (Abcam), G6pdx (Abcam), *α*-tubulin (Santa Cruz).

### QUANTIFICATION AND STATISTICAL ANALYSIS

#### Bulk RNAseq analysis

Raw reads from bulk RNA-seq were aligned to the mm10 genome using STAR v 2.6.1d with the quantmode parameter to get read counts. DESeq2 v 1.22.2 was used for differential expression analysis. Heatmaps were produced using gplots v 3.0.1.1.

#### Pre-processing of scRNA-seq data

The single cell RNA raw sequence reads were aligned to the UCSC Mus musculus genome, mm10, using Kallisto (Bray et al., 2016). The mean number of reads per cell was 241,500. Gene expression values were quantified into Transcripts Per Kilobase Million (TPM). We removed cells which had fewer than 800 detected genes or more than 3,000 genes. After filtering, the mean number of detected genes per cell were 1,765. Then, for the SFB and EAE dataset, we normalized the gene matrix by SCTransform (Hafemeister and Satija, 2019) regression with percent of mitochondria reads separately.

#### Visualization and clustering

To visualize the EAE and SFB data, we performed principal component analysis (PCA) for dimensionality reduction, and further reduced PCA dimensions into two dimensional space using UMAP (Becht et al., 2018; McInnes and Healy, 2018). UMAP matrix was also used to build the weighted shared nearest neighbor graph and clustering of single cells was performed by the original Louvain algorithm. All the above analysis was performed using Seurat v3 R package (Butler et al., 2018; Stuart et al., 2019). However, we noticed there was one cluster of cells with a much lower number of features in which over 60% of cells had fewer features than the average. Thus, it was removed from the analysis.

A table of average gene expression for each cluster, and a table of log2 Fold Change values with p-values between each sample were calculated with Seurat. Seurat calculates differential expression based on the non-parameteric Wilcoxon rank sum test. Log fold change is calculated using the average expression between two groups. These values were used in Figure S1 J. and K. with IPA for a pathway analysis, and in Figure S1 L. to make volcano plots.

#### Statistical analysis

Differences between groups were calculated using the unpaired two-sided Welch’s t-test. Difference between two groups of co-transferred cells, or in bone marrow chimera experiments, were calculated using the paired two-tailed t-test. Differences between EAE clinical score was calculated using two-way ANOVA for repeated measures with Bonferroni correction. For RNA-seq analysis, differential expression was tested by DESeq2 by use of a negative binomial generalized linear model. We treated less than 0.05 of p value as significant differences. ∗p < 0.05, ∗∗p < 0.01, ∗∗∗p < 0.001, and ∗∗∗∗p < 0.0001. Details regarding number of replicates and the definition of center/error bars can be found in figure legends.

